# Drought stress increases the expression of barley defence genes with negative consequences for infesting cereal aphids

**DOI:** 10.1101/2021.09.12.459767

**Authors:** Daniel J. Leybourne, Tracy A Valentine, Kirsty Binnie, Anna Taylor, Alison J Karley, Jorunn IB Bos

## Abstract

Crops are exposed to myriad abiotic and biotic stressors with negative consequences. Two stressors that are expected to increase under climate change are drought and infestation with herbivorous insects, including important aphid species. Expanding our understanding of the impact drought has on the plant-aphid relationship will become increasingly important under future climate scenarios. Here we use a previously characterised plant-aphid system comprising a susceptible variety of barley, a wild relative of barley with partial-aphid resistance, and the bird cherry-oat aphid to examine the drought-plant-aphid relationship. We show that drought has a negative effect on plant physiology and aphid fitness and provide evidence to suggest that plant resistance influences aphid responses to drought stress, with the expression of aphid detoxification genes increasing under drought when feeding on the susceptible plant but decreasing on the partially-resistant plant. Furthermore, we show that the expression of thionin genes, plant defensive compounds that contribute aphid resistance, increase ten-fold in susceptible plants exposed to drought stress but remain at constant levels in the partially-resistant plant, suggesting they play an important role in modulating aphid populations. This study highlights the role of plant defensive processes in mediating the interactions between the environment, plants, and herbivorous insects.

## Introduction

Under the changing climate it is anticipated that annual levels of precipitation will decrease in some regions, resulting in extended periods of global drought (Blenkinsop and Fowler 2007; Santos et al., 2016). Exposure to drought can have severe impacts on plant physiology, often leading to reduced growth and photosynthetic capacity, leading to reductions in crop yields (Osakabe et al., 2014; Zeppel et al., 2014). Drought stress can have wider impacts on the relationships between plants and insects. These consequences are primarily mediated through plant physiological (Osakabe et al., 2014; Zeppel et al., 2014) and molecular (Ozturk et al., 2002; Davila Olivas et al., 2016) responses to drought stress, and the associated changes can also influence the population dynamics, fitness, phenology, biology, and behaviour of herbivorous insects (Huberty and Denno, 2004; Mody et al., 2009; Aslam et al., 2013; Kansman et al., 2020; Lin et al., 2021). The responses of herbivorous insects to drought stressed plants can vary widely (Pons et al., 2020; Xie et al., 2020; Cui et al., 2021) and the underlying reasons for these varying responses are not always clear.

Drought stress is primarily associated with a reduction in soil water content and the availability of water for uptake by the plant roots. Soil physical properties, including pore size, soil strength, aeration, and bulk density, can also influence the structure of the root system and impact the effectiveness of water uptake (Bengough et al., 2006; Bengough et al., 2011; Haling et al., 2014; Valentine et al., 2012). For example, under drought there is a general increase in soil strength, which increases the energy required for root elongation, altering the structure of the root system and decreasing water accessibility (Colombi et al., 2018). Soil density also affects water availability under drought: in loose soils, root-soil contact can be reduced, increasing the severity of drought effects, whereas in more compacted soils, root-soil contact can increase, which improves water uptake even when root elongation is reduced. Soil pore size can influence the extent to which water contained within the pores is accessible (water contained in smaller pores is harder to access than water contained in larger pores). Therefore, changes in soil strength, compaction, or porosity can initiate, or exacerbate, the level of drought stress experienced by plants (Schmidt et al., 2012).

Physiological responses of plants to drought stress generally include reduced cell turgor, stomatal pore closure, reduced growth and productivity, reduced water use efficiency, and a decreased rate of photosynthesis (Osakabe et al., 2014; Zeppel et al., 2014). Drought-mediated changes in plant physiology are widely reported (Cornelissen et al., 2008) and have been associated with negative consequences for herbivorous insects across multiple herbivorous insect groups (Huberty & Denno 2004). A recent meta-analysis of aphid responses to drought stress in plants indicated that drought-induced reduction in aphid fitness is also associated with a reduction in plant growth and vigour and an increase in the tissue concentrations of plant defensive compounds (Leybourne et al., 2021). There is evidence that plant resistance against aphids can mediate the extent to which aphids are negatively affected by drought, with aphids feeding on resistant plants showing a smaller decrease in fitness under drought conditions compared with aphids feeding on susceptible plants (Oswald and Brewer, 1997; Dardeau et al., 2015). Our recent meta-analysis highlighted a knowledge gap regarding the interaction between plant resistance traits and drought (Leybourne et al., 2021). Examining these interactive effects is becoming more important as alternative pest management methods (including breeding for plant resistance against insect pests) are increasingly needed and droughts are more likely in the future.

Barley is an important crop for the UK economy, the fourth most important cereal crop worldwide (Newman and Newman, 2006; Newton et al., 2011), and the barley industry is extremely vulnerable to drought (Xie et al., 2018). A further factor which negatively affects crop yields is biotic stress introduced by pests and pathogens (Perry et al., 2000). For example, infestation with aphids, a phloem feeding herbivorous insects with a global distribution (Blackman and Eastop, 2000), can lead to substantial decreases in crop yields. Crop losses can be further exacerbated by the transmission of aphid-vectored plant viruses (Smith and Sward, 1982; Perry et al., 2000; Murray and Brennan, 2010). Under a changing climate herbivorous insects are anticipated to become an increasingly significant cause of biotic stress to agricultural crops (Deutsch et al., 2018). The wild progenitors of those crop species are an important source of agronomic traits that, when exploited, could improve the tolerance of modern crops to abiotic stress and increase crop resistance against insect herbivores and plant pathogens (Ellis et al., 2000; Dempewolf et al., 2014).

Defences involved in aphid resistance in *Hordeum spp*. include increased expression of thionin (antimicrobial peptides) genes (Delp et al., 2009; Mehrabi et al., 2014; Escudero-Martinez et al., 2017), increased chitinase and *β*-1,3-glucanase activity (Forslund et al., 2000), and the presence of plant secondary metabolites (Gianoli and Niemeyer, 1998). Plant phytohormone signalling pathways, including Abscisic Acid (ABA), Salicylic Acid (SA), Jasmonic Acid (JA) and Ethylene (ET) signalling, mediate coordinated molecular responses to herbivory via the regulation of defence signalling genes and the biosynthesis of defensive allelochemicals (Smith and Boyko, 2007; Bari and Jones, 2009; Morkunas et al., 2011; Foyer et al., 2016); higher constitutive expression of phytohormone signalling genes can lead to improved resistance against aphids in cereals (Losvik et al., 2017). A wild progenitor species of barley, Hsp5 (*H. spontaneum* 5), has partial-resistance against two important cereal aphid species, the bird cherry-oat aphid (*Rhopalosiphum padi*) and the grain aphid (*Sitobion avenae*) (Delp et al., 2009; Leybourne et al., 2019). Partial-resistance against aphids in Hsp5 is associated with increased expression of thionin and phytohormone (abscisic acid and ethylene) genes alongside a decrease in the concentration of essential amino acids in plant phloem, when compared with susceptible modern barley cultivars (Leybourne et al., 2019). Physiological and biochemical investment in constitutive defence against aphids may compromise the ability of plants to respond to abiotic stress, resulting in a potential trade-off in resource allocation under dual biotic and abiotic stress (Lin et al., 2021).

Aphid infestation of plants is facilitated by the secretion of effector molecules into plant tissue. Effectors are small molecules contained in aphid saliva that are secreted during aphid probing where they bind with plant proteins to suppress plant defences (as reviewed by Hogenhout & Bos, 2011). Overexpression of effector genes *Rp1* and *RpC002* from *R. padi* in barley increased barley susceptibility to *R. padi* (Escudero-Martinez et al., 2020). Interestingly, transgenic barley lines expressing *Rp1* showed reduced defence gene expression, including reduced expression of a defensive *β*-*thionin* gene (Escudero-Martinez et al., 2020). Further, aphids deploy a suite of molecular mechanisms to detoxify plant defensive biochemical compounds that might be ingested during aphid feeding (Pontoppidan et al., 2001; Francis et al., 2002), resulting in a dynamic molecular landscape at the plant-aphid interface. How an abiotic stress (such as drought stress) might impact on these molecular interactions is unclear. For example, increased expression of plant defensive processes under drought (Ozturk et al., 2002) might depend on the level of constitutive defence against aphids (Leybourne et al., 2019), leading to differential molecular responses of aphids to drought on aphid-resistant and -susceptible plants,

In this study, the physiological and molecular interactions between barley (*Hordeum vulgare* cv. Concerto) and a wild progenitor species of barley, Hsp5 (*H. spontaneum* 5), with the bird cherry-oat aphid (*Rhopalosiphum padi*) were examined under control and drought stress conditions. We developed two drought stress treatments using a calibration curve devised from excavated field soil, one of which was designed to impose an additional root impedance treatment through higher soil compaction (Valentine et al., 2012). We use this study-system to examine the effects of plant drought stress on plant physiology and defence, and aphid fitness and feeding/detoxification processes.

## Materials & Methods

### Development of drought stress treatments, aphid rearing conditions, and seed germination conditions

The development of the drought stress treatments is described in detail in Supplementary File 1. Briefly, field soil was excavated, sieved, compacted into circular cores at two densities (1.15 g cm^3^ and 1.25 g cm^3^) and subjected to suction at a range of matric potentials to establish water release curves, to assess soil strength, and to quantify soil physical characteristics (see Valentine et al., 2012). Three water treatments were designed based on the soil water retention and soil strength data: i) a control treatment (*c*. 40 % gravimetric moisture content (gMC); 1.15 g cm^3^ soil dry bulk density, with soil strength estimated to be 1.06 MPa, and an estimated matric potential of 15-20 -kPa) which was below field capacity but with an acceptable level of aeration and a soil strength which would not impede root growth; ii) a water-limited treatment (*c*. 30 % gMC; 1.15 g cm^3^ dry bulk density, *c*. 350 -kPa matric potential) which had a lower gravimetric moisture content than the control treatment but remained above the permanent wilting point with a soil strength that was higher than the control but only partially limiting to root growth (soil strength estimated to be 2.24 MPa); and iii) a combined stress treatment (*c*. 30 % gMC; 1.25 g cm^3^ dry bulk density, soil strength estimated to be 3.92 MPa, *c*. 400-500 -kPa matric potential) which had a lower gravimetric moisture content than the control treatment but remained above the permanent wilting point with a soil strength which should impede root growth significantly.

Asexual laboratory cultures of the bird cherry-oat aphid, *Rhopalosiphum padi* Linnaeus were established from individual apterous adults collected from the James Hutton Institute, Dundee, UK (characterised in Leybourne et al. (2020)). Aphid cultures were reared on one-week old barley seedlings (cv. Optic) contained in ventilated cups at 20 °C and 16:8 h (L:D). *Hordeum vulgare* Linnaeus cv. Concerto (Concerto) and *H. spontaneum* 5 Linnaeus (Hsp5) seeds were surface-sterilised by washing in 2% (v/v) hypochlorite and rinsing with d.H_2_O. Seeds were kept moist and in the dark: Hsp5 seeds were incubated at 4°C for 14 days (to break dormancy) and Concerto seeds were kept at room temperature for 48 h; all seedlings were at a similar developmental stage (GS07) when transferred to the soil. Germinated seedlings were planted into soil collected from the Mid Pilmore field at the James Hutton Institute, Invergowrie, Dundee, UK under one of the three water-stress treatments described above and in Supplementary File 1 and grown in randomised blocks under controlled glasshouse conditions with a 16:8 h day:night length and a 20:15°C day:night temperature. Fig. 1A shows the relative tensiometer measurements of the soil matrix over the lifetime of the experiments.

**Fig. 1:**
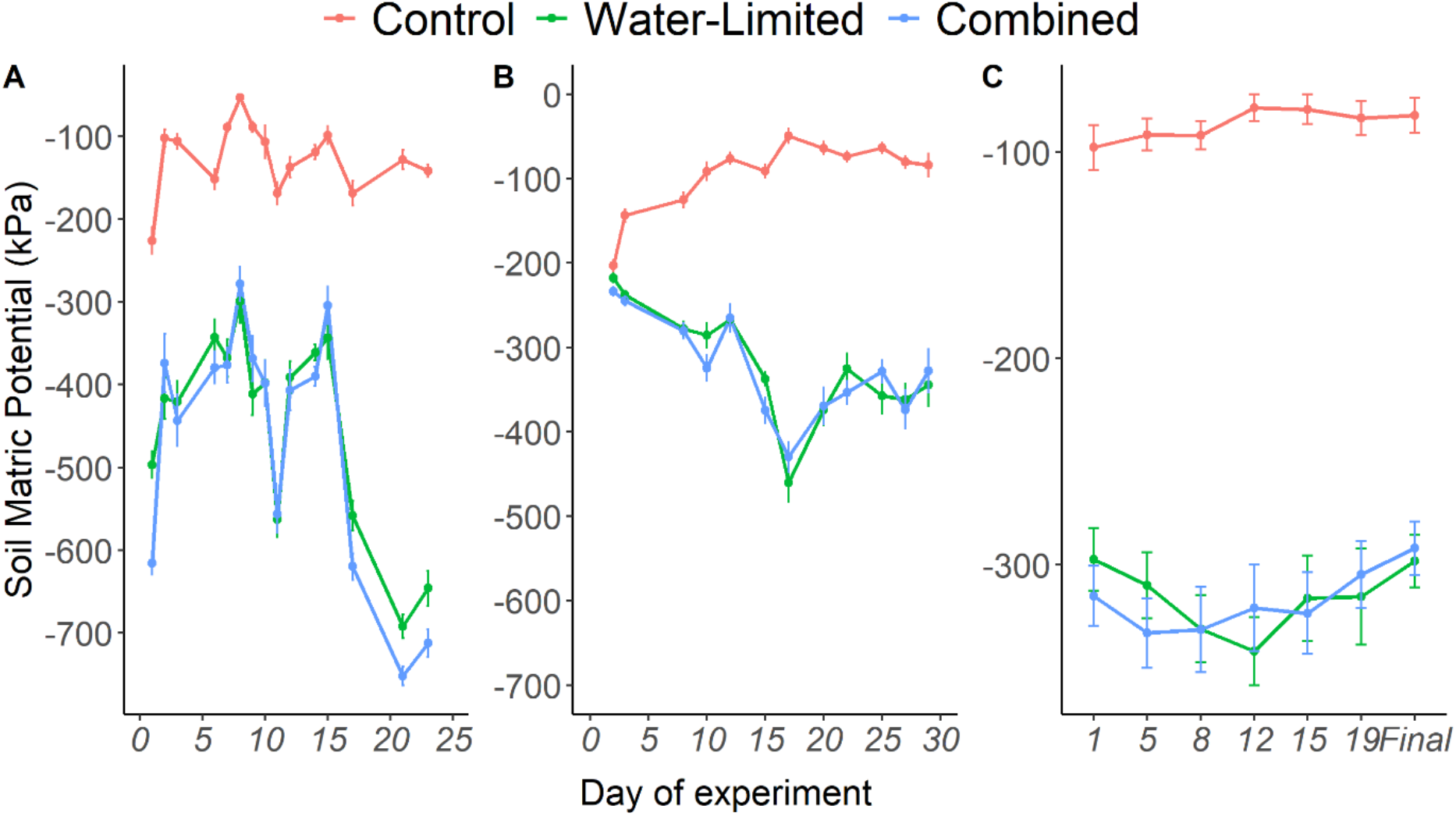
Matric potential (kPa: mean ± se) of the soil matrix over the course of the experiments. A) Matric potential in temporal block A of the main experiment; B) Matric potential in temporal block B of the main experiment; C) Matric potential during plant growth prior to use in the EPG experiment alongside the final reading taken immediately prior to EPG analysis.

### Experimental design of drought stress study

Germinated seedlings were grown in Perspex cylindrical tubes (internal volume *c*. 950 cm^3^; internal diameter *c*. 5 cm) under one of the three water treatments described in Supplementary File 1. Plants were grown in fully randomised blocks in a split-plot experimental design incorporating two temporal whole plots and ten blocks within each temporal whole plot, with each block containing one replicate of each of twelve treatment combinations: two plant types (Hsp5, Concerto), three soil treatments (control, water-limited, combined) and two aphid treatments (empty clip-cages (MacGillivray and Anderson, 1957) or clip-cages containing three adult apterous *R. padi* (Genotype G; Leybourne et al., 2020). Plants were grown under the desired soil treatment until the first true-leaf developmental stage (GS12), at which point the aphid treatment was introduced.

Plants were monitored daily to record development time until the first true-leaf stage; stomatal aperture was measured at three time-points with readings taken from the tip of the flag leaf. When all plants reached the first true-leaf stage the aphid treatments (aphid-free clip-cage control, or clip-cage containing three adult apterous aphids) were introduced to each plant. Two experimental harvests were carried out: a subset, *n =* 6 (3 per each temporal whole plot), was collected 24 h post-aphid infestation for analysis of plant and aphid molecular responses, and a larger subset, *n = 14* (7 per each temporal whole plot), was collected 7-days after aphid infestation for plant physiological and aphid fitness responses. Blocks were selected semi-randomly for the 24 h harvest (ensuring that at least one block was taken from the East and West side of the glasshouse, representing the environmental gradient in the cubicle).

Plants were harvested by cutting the stem at the stem-root junction, excising the first true leaf (leaf with clip-cage attached) from the plant stem and further separating the leaf material inside the clip-cage from the rest of the true-leaf. If present, aphids were removed, and the plant tissue contained within the clip-cage was weighed and immediately flash-frozen in liquid nitrogen. Adult aphids were then collected and immediately flash-frozen in liquid nitrogen, the aphid nymphs were counted and then flash-frozen in liquid nitrogen, freeze-dried, and weighed. The remainder of the true-leaf and the remaining above-ground plant tissue were weighed and all material was flash-frozen in liquid nitrogen. All flash-frozen material was stored at −80°C: leaf material from inside the clip-cage and adult aphids were stored until RNA extraction and all other material was freeze-dried and weighed. After collecting aphid and plant shoot tissue, the soil cores were removed from the Perspex tubes and plant roots were carefully removed from the soil matrix for recording root length (measuring from the root-stem junction to the end of the root system using a ruler). Roots were washed and dried at 70°C for 48h, and root dry mass was recorded; root:shoot allometry was calculated using root dry mass and above-ground freeze-dried mass.

### Plant and aphid gene expression analysis

RNA was extracted from collected plant and aphid tissue from the first experimental harvest (24 h after aphid infestation) by grinding collected tissue to a fine powder using a mortar and pestle under liquid nitrogen. Total RNA was extracted using the Norgen Plant-Fungi RNA Extraction kit, following the manufacturer’s protocol and including on-column DNAse treatment. cDNA synthesis was carried out using the SuperScript III™ cDNA synthesis kit following the manufacturer’s protocol: for aphids, approximately 250 ng RNA was used in the cDNA synthesis reactions, with *c*. 1000 ng used in barley cDNA synthesis. RT-qPCR primers (Table S1 and Table S2) were designed using the Universal Probe Library Assay Design Centre (Roche Life Sciences) and primer efficiency was determined using pooled cDNA samples. The GeNorm procedure (Vandesompele et al., 2002) was used to identify suitable reference genes with stable expression levels under all experimental conditions. For Concerto and Hsp5 these were *HvGR* (Barley gene index - HvGI: TC146685) and *HvEF*-*1*-*α* (HvGI: TC146566) (Table S1); for *R. padi* only one reference gene was determined to be sufficiently stable, *RpCDC42* (Table S2). RT-qPCR assays were carried out in triplicate using SYBR® Green chemistry with GoTaq® qPCR Master Mix (Promega, UK) on a StepOne™ Real-Time PCR Machine (Applied Biosystems, UK). Reactions were carried out in a final volume of 12.5 μl with a final concentration of 1x GoTaq® qPCR Master Mix, 1 μM of each primer, 1.4 mM MgCl_2_, 2.4 μM CXR reference dye and approx. 12.5 ng of plant cDNA or 4 ng of aphid cDNA. RT-qPCR conditions were as follows: 95°C for 15 mins followed by 40 cycles of denaturing for 15 s at 95°C, annealing for 30 s at 60°C and 30 s at 72°C for DNA extension. Fluorescence was recorded at the end of each annealing cycle and a melting curve was incorporated into the end of the RT-qPCR programme. The 2^−ΔΔCt^ method (Livak and Schmittgen, 2001) was used to calculate values relative to the mean of the reference genes. Values were further normalised to the internal control within each block (the Concerto-control-no aphid treatment for the plant samples, and Concerto control treatment for the aphid samples).

### Experimental design of aphid feeding behaviour measurements

Hsp5 and Concerto plants were grown in Perspex cylindrical tubes (internal volume *c*. 317.5 cm^3^; internal diameter *c*. 2.5 cm) under one of the three water treatments described above (see Fig. 1C for the soil matrix water levels for these plants). Plants were grown until the first true-leaf development stage (GS12) at which point they were used in aphid feeding assays. The DC-EPG technique (Tjallingii, 1978; Tjallingii, 1988; Tjallingii, 1991; Tjallingii, 2001) was employed to monitor the probing and feeding behaviour of adult apterous *R. padi* over a six hour period using a Giga-4 DC-EPG device (EPG Systems, The Netherlands). The order in which plant – drought combinations were tested and allocated to each of the three EPG probes was randomised. Data were acquired using Stylet+D software (EPG Systems, The Netherlands). Aphids were lowered onto the first true-leaf immediately after the recording started. Successful recordings were made for each plant-soil treatment combination as follows: nine (Concerto control), seven (Concerto water-limited), eight (Concerto combined), ten (Hsp5 control), seven (Hsp5 water-limited), and 11 (Hsp5 combined). All EPG recordings were obtained within a grounded Faraday cage.

EPG waveforms were annotated using Stylet+A software (EPG Systems, The Netherlands). Waveforms were annotated by assigning waveforms to np (non-probing), C (stylet penetration/pathway), pd (potential-drop/intercellular punctures), E1 (saliva secretion into phloem), E2 (saliva secretion and passive phloem ingestion), F (penetration difficulty) or G (xylem ingestion) phases (Tjallingii, 1988; Alvarez et al., 2006). No E1e (extracellular saliva secretion) phases were detected. Annotated waveforms were converted into time-series data using the excel macro developed by Schliephake et al., 2013 (Julius Kühn-Institut, Germany; available at www.epgsystems.eu).

### Statistical analysis

All statistical analyses were carried out using R Studio Desktop version 1.0.143 running R version 3.5.1 (R-Core 2019), with additional packages ade4 v.1.7-16 (Dray & Dufour, 2007), car v.2.1-4 (Fox and Weisberg, 2011), eha v.2.5.1 (Broström, 2012), emmeans v.1.3.1 (Lenth, 2018), factoextra v1.0.7, ggplot2 v.2.2.1 (Wickham, 2009), ggpubr v.0.1.2 (Kassambara, 2017), ggfortify v.0.4.5 (Tang et al., 2016), lme4 v.1.1-13 (Bates et al., 2015), lmerTest v.2.0-33 (Kuznetsova et al., 2017), lsmeans v.2.27-62 (Lenth, 2016), nlme v.3.1-131, Multcomp v.1.4-8 (Hothorn et al., 2008), pkbrtest v.0.4-7 (Halekoh and Højsgaard, 2014), and vegan v.2.5-7.

#### Multivariate redundancy analysis of plant physiology, aphid fitness, and EPG data

Redundancy analysis was used to determine how drought stress affects plant physiology (for individual parameter results see Table S3 and Table S6 for the statistical results) and aphid fitness (for complete results of aphid fitness and aphid feeding experiments see Tables S4; S7, respectively, and Table S5 for the statistical results). Plant and aphid responses to drought stress were pooled into three multivariate datasets: plant data from the primary drought stress experiment (shoot dry mass, root dry mass, root length, root:shoot allometry, average stomatal conductance, and plant development time to the true leaf stage), aphid fitness data from the primary drought stress experiment (comprising dry mass of nymphs and cumulative adult fecundity over seven days), and aphid feeding data from the supplementary EPG experiment (all 103 EPG variables assessed). First, multivariate analysis was carried out using principal components analysis, testing each multi-variate dataset against all possible explanatory variables (for plant data this included plant type, drought treatment, aphid infestation, and all interactions; for aphid data explanatory variables included plant type, drought treatment, and the interaction). Models were simplified through manual backward model selection and Permutated Analysis of Variance was carried out on the final redundancy analysis model for each multi-variate dataset to identify which explanatory variables significantly influenced the observations.

#### Standard analysis of plant physiology, aphid fitness, and EPG data

All plant physiology parameters were modelled as response variables against drought treatment, plant type, aphid presence (if applicable) and the interaction; all aphid data (fitness and EPG data) were modelled as response variables against drought treatment, plant type, and the interaction. Plant development time was modelled using event history analysis (survival analysis) by fitting a cox mixed effects regression model, with temporal block and sub-block incorporated as random factors. A X^2^ test was carried out on the final model. Stomatal conductance was analysed in a mixed effects model fitted with a temporal correlation structure for repeated measures analysis. Plant shoot dry mass, root dry mass, root length, root:shoot allometry, aphid dry mass, and aphid fecundity were analysed in separate linear mixed effects models, with temporal block and sub-block included as random factors. Type II Analysis of Variance tests were applied to the final models. EPG data were analysed by fitting a permutated multiple analysis of variance to the entire EPG dataset.

#### Analysis of RT-qPCR data

Plant and aphid gene expression data (the calculated 2^−ΔΔCt^ value) were analysed with generalised least square estimation models. For plant genes, each gene was modelled in response to plant type, water treatment, aphid presence, and all interactions. For aphid data, genes were modelled in response to plant type, water treatment, and the interaction. Models were weighted for variance introduced between the experimental blocks by accounting for fixed variation between the blocks (using the “varFixed” function) and incorporating this into the models using the “weights” function. Final models were analysed with a χ^2^ test and fitted-residual plots were used to assess model suitability. *HvTHIO*, *HvA1*, *RpCOO2*, *RpSec27*, *RpSec22*, *RpSec5*, *RpAD*, *RpMYR*, and *RpGG* were fitted with a log transformation; *HvERF1*, *HvNPR1 and RpMAPK4* were fitted with a sqrt transformation; *HvLOX2* was not transformed. General linear hypothesis testing with single-step p-value adjustment was used as a *post*-*hoc* test for each model. Gene and primer information can be found in Table S1 and Table S2.

## Results

### Drought stress has an adverse effect on plant physiology, aphid fitness, and aphid feeding behaviour

A range of plant physiological, aphid fitness, and aphid feeding parameters were measured under control, water-limited, and combined stress treatments (See Tables S3 – S5 for full results and Tables S6 – S7 for the statistical results for each individual parameter). Multi-variate analysis of these parameters indicated that drought stress treatment significantly affected plant physiology (F_2,163_ = 35.28; p = 0.001; Fig. 2A), aphid fitness (F_2,81_ = 6.51; p = 0.002; Fig. 2B), and aphid feeding behaviour (F_2,49_ = 4.99; p = 0.001; Fig. 2C). Redundancy analysis showed that significant variation in response to the different drought treatments was explained by the first dimension for plant physiological data (F_1,163_ = 77.10; p = 0.001), aphid fitness data (F_1,81_ = 13.03; p = 0.002), and aphid feeding data (F_1,49_ = 8.27; p = 0.001), but other dimensions did not vary significantly in relation to these treatment factors. Differences in plant physiology were also detected between the two plant types (F_1,163_ = 8.08; p = 0.001) and in response to aphid infestation (F_1,163_ = 4.39; p = 0.025), however no interactive treatment effects were identified for plant physiological parameters. Similarly, no differences in aphid fitness or aphid feeding behaviour were detected between the two plant types or in response to the drought stress x plant interaction.

**Fig. 2:**
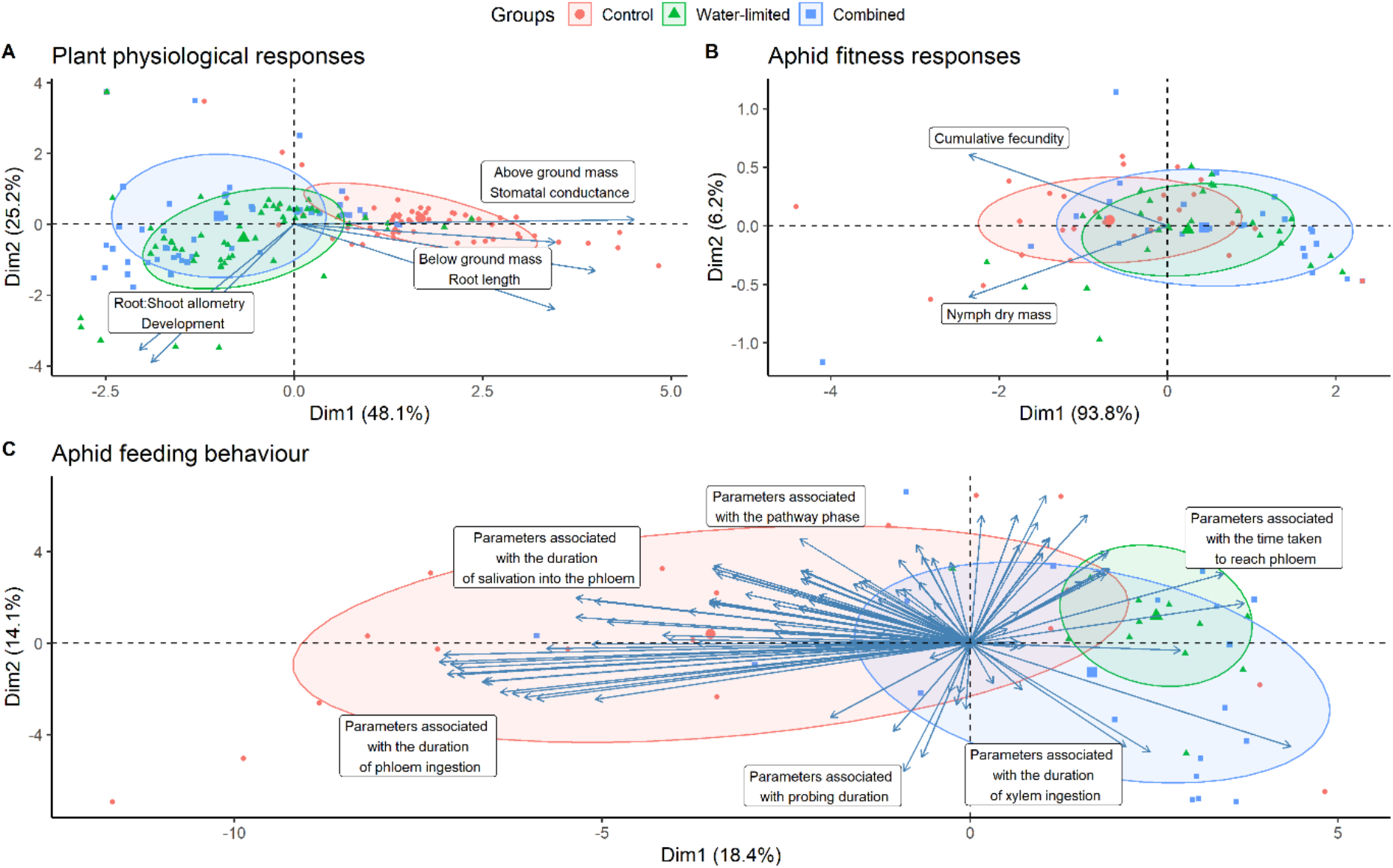
The first two dimensions of multi-dimensional scaling models for all variables describing (A) plant physiology (shoot dry mass, root dry mass, root length, root:shoot allometry, average stomatal conductance, and plant development time to the true leaf stage), (B) aphid fitness (comprising dry mass of nymphs, and cumulative fecundity over seven days), and (C) aphid feeding (all 98 EPG variables assessed) in response to the applied water treatments: control, water-limiting, and combined. Axes show the first two dimensions and the variation (%) explained by each dimension.

The effect of drought stress on aphids included a 30% reduction in the cumulative fecundity for aphids feeding on both plant types under drought conditions compared with the fecundity of aphids under control conditions (Table S4; S7), with similar levels of reduction observed for aphid mass. The negative effect of drought on aphid feeding behaviour (Table S5; S7) included reduced phloem ingestion and increased xylem ingestion for aphids exposed to drought stress conditions compared with aphids under control conditions.

The plant physiological parameters affected by drought stress (Table S3; S6) included development time (slower under both drought stress conditions imposed), above ground and below ground dry mass (lower for both parameters under both drought stress conditions imposed, compared with control conditions), and root:shoot allometry (higher for plants exposed to both drought stress conditions compared with control conditions).

A further physiological parameter affected by drought stress was stomatal conductance. Plants exposed to drought stress (water-limited and combined treatments) had a smaller stomatal conductance compared with plants under control conditions; additionally, the stomatal conductance of Hsp5 was consistently smaller than that of Concerto under all three treatments (Table S3). Root length also differed between the two plant types, although no other physiological differences between Hsp5 and Concerto were identified.

### HvTHIO1 gene expression is elevated in Concerto under drought stress conditions and HvA1 is more highly expressed in Hsp5 under all treatments

The drought treatments had a significant effect on the expression of three of the barley defence-related genes assessed, *HvTHIO1*, *HvA1*, *HvERF1* (Table 1; Fig. 3). Selection of these defence genes was based on their association with Hsp5 partial resistance against aphids (Leybourne et al., 2019). *HvTHIO1* expression levels were higher in Hsp5 compared with Concerto under control conditions. However, the two plant types responded differentially to drought stress: *HvTHIO1* expression increased in Concerto under drought (Table 1; Fig. 3A), such that it was significantly higher in Concerto than Hsp5 in the water-limited and combined stress treatments.

**Table 1:** Statistical results of generalised least squares estimation models for plant gene expression results in response to plant, water treatment, aphid infestation, and all interactions. Bold text indicates significant p values.

| Explanatory variable | Gene |  |  |  |  |  |  |  |  |  |
| --- | --- | --- | --- | --- | --- | --- | --- | --- | --- | --- |
|  | *HvTHIO1* |  | *HvA1* |  | *HvERF1* |  | *HvLOX2* |  | *HvNPR1* |  |
| Plant Type | $X^2_1 = 1.01$ | $p = 0.313$ | $X^2_1 = 16.63$ | **$p < 0.001$** | $X^2_1 = 3.67$ | $p = 0.055$ | $X^2_1 = 14.90$ | **$p < 0.001$** | $X^2_1 = 2.34$ | $p = 0.125$ |
| Water Treatment | $X^2_2 = 6.54$ | **$p = 0.037$** | $X^2_2 = 10.28$ | **$p = 0.005$** | $X^2_2 = 9.82$ | **$p = 0.007$** | $X^2_2 = 1.64$ | $p = 0.558$ | $X^2_2 = 4.91$ | $p = 0.083$ |
| Aphid Presence | $X^2_1 = 0.08$ | $p = 0.771$ | $X^2_1 = 0.15$ | $p = 0.695$ | $X^2_1 = 0.01$ | $p = 0.907$ | $X^2_1 = 11.52$ | **$p < 0.001$** | $X^2_1 = 0.92$ | $p = 0.335$ |
| Experimental Block | $X^2_1 = 0.01$ | $p = 0.893$ | $X^2_1 = 9.09$ | **$p = 0.002$** | $X^2_1 = 1.89$ | $p = 0.168$ | $X^2_1 = 0.97$ | $p = 0.323$ | $X^2_1 = 0.74$ | $p = 0.386$ |
| Plant x Water Treatment | $X^2_2 = 15.53$ | **$p < 0.001$** | $X^2_2 = 1.84$ | $p = 0.397$ | $X^2_2 = 0.84$ | $p = 0.655$ | $X^2_2 = 3.94$ | $p = 0.139$ | $X^2_2 = 2.78$ | $p = 0.249$ |
| Plant Type x Aphid Presence | $X^2_1 = 2.31$ | $p = 0.127$ | $X^2_1 = 0.57$ | $p = 0.447$ | $X^2_1 = 0.01$ | $p = 0.964$ | $X^2_1 = 0.14$ | $p = 0.707$ | $X^2_1 = 1.30$ | $p = 0.252$ |
| Water Treatment x Aphid Presence | $X^2_2 = 3.80$ | $p = 0.149$ | $X^2_2 = 2.02$ | $p = 0.362$ | $X^2_2 = 0.56$ | $p = 0.742$ | $X^2_2 = 2.64$ | $p = 0.266$ | $X^2_2 = 0.48$ | $p = 0.783$ |
| Plant Type x Water Treatment x Aphid Presence | $X^2_2 = 1.02$ | $p = 0.599$ | $X^2_2 = 0.31$ | $p = 0.853$ | $X^2_2 = 0.66$ | $p = 0.716$ | $X^2_2 = 1.43$ | $p = 0.487$ | $X^2_2 = 0.91$ | $p = 0.632$ |

**Fig. 3:**
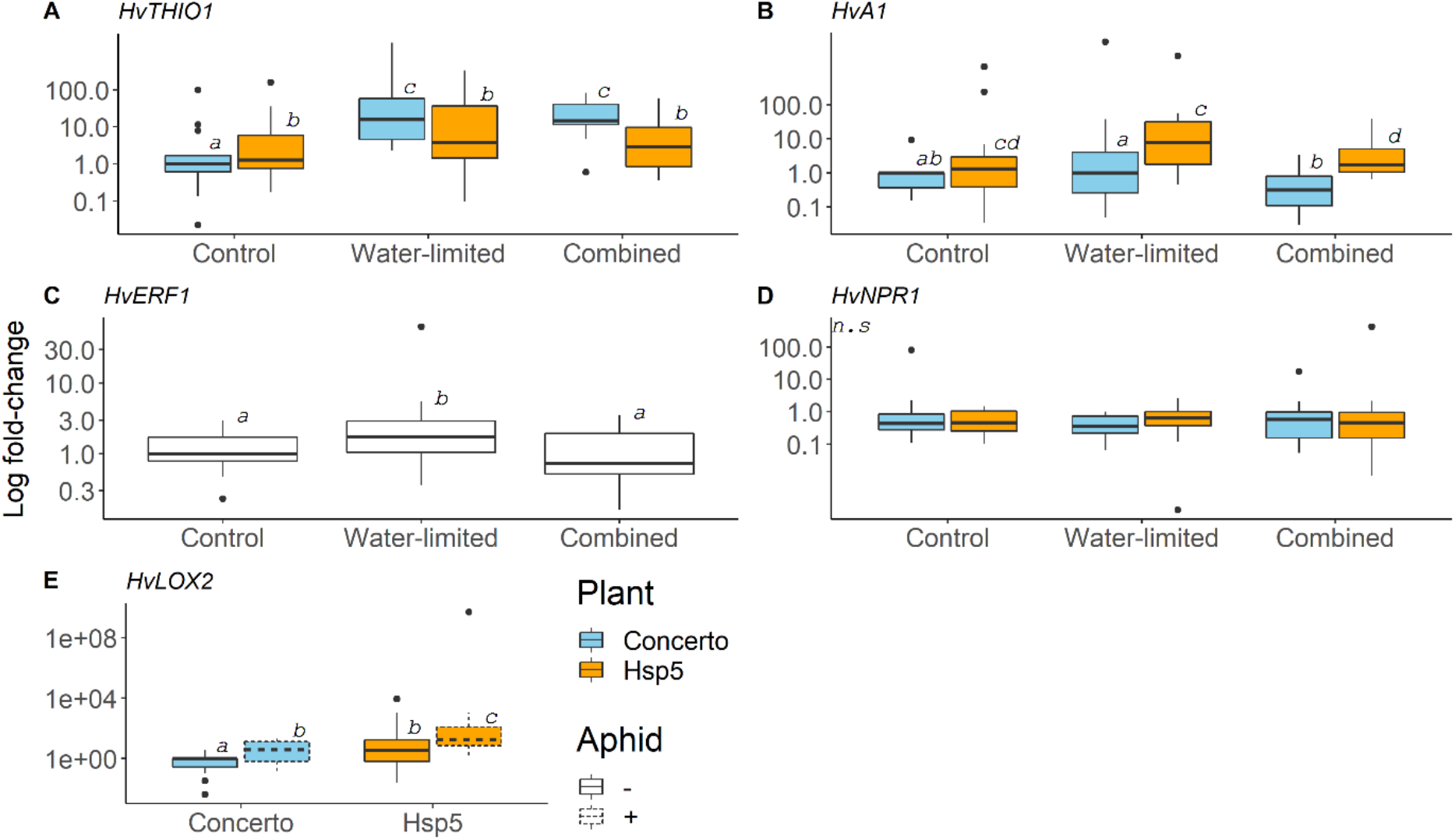
Gene transcript levels of (A) *HvTHIO1* (defensive thionin), (B) HvA1 (ABA-responsive), (C) *HvERF1* (ET-responsive), (D) *HvNPR1* (SA-responsive) and (E) *HvLOX2* (JA-responsive). *HvTHIO1* (n = 12), HvA1 (n = 12) and *HvNPR1* (n = 12) are shown for both plant types in response to the different water treatments: control, water-limited, and combined stress (Combined). Expression levels of *HvERF1* (n = 24) are shown in response to the different water stress treatments only. . *HvLOX2* (n = 18) expression is shown in response to aphid treatment (solid line – no aphid; dashed line – aphid infestation for 24 h) for both plant types. All gene expression values are relative to the mean expression of two reference genes, *HvGR* and *HvEF*-*1*-*α*, and data are presented on the log_10_ scale. Letters show significant differences based on general linear hypothesis testing with single-step p-value adjustment post-hoc analysis.

*HvA1* expression levels were higher in Hsp5 compared with Concerto under all treatments but were reduced under the combined stress treatment compared with the water-limited treatment in both plant types (Table 1; Fig. 3B). *HvERF1* expression levels were higher under water-limited conditions compared with control and combined conditions for both plant types (Table 1; Fig. 3C). *HvNPR1* expression was not differentially affected by any variable assessed (Table 1; Fig. 3D) and *HvLOX2* was differentially expressed between the two plant types (higher in Hsp5) and increased in response to aphid infestation (Table 1; Fig. 3E).

### The expression of aphid detoxification genes is differentially affected by host plant and drought stress

To investigate whether drought stress affected aphid effector and detoxification gene expression, we analysed expression levels of the *R. padi* effectors *RpC002*, *RpSec5*, *RpSec22*, *RpSec27*, the stress-responsive genes *RpMAPK4* and *RpAD*, and detoxification genes *RpGG* and *RpMyr* using qRT-PCR. The selected *R. padi* effectors and *RpMAPK4* were not affected by host plant type, water treatment or a plant type x water treatment interaction (Table 2; Fig. 4).

**Table 2:** Statistical results of aphid gene expression analysis. Bold text indicates significant p values.

| Gene | Explanatory Variable |  |  |  |  |  |  |  |
| --- | --- | --- | --- | --- | --- | --- | --- | --- |
|  | Plant Type |  | Water Treatment |  | Experimental Block |  | Plant Type x Water Treatment |  |
| *RpC002* | $X^2_1 = 0.01$ | $p = 0.896$ | $X^2_2 = 4.06$ | $p = 0.130$ | $X^2_1 = 6.37$ | $p = \mathbf{0.011}$ | $X^2_2 = 3.32$ | $p = 0.189$ |
| *RpSec5* | $X^2_1 = 3.36$ | $p = 0.066$ | $X^2_2 = 1.38$ | $p = 0.500$ | $X^2_1 = 20.69$ | $p = \mathbf{0.001}$ | $X^2_2 = 1.37$ | $p = 0.500$ |
| *RpSec22* | $X^2_1 = 3.32$ | $p = 0.068$ | $X^2_2 = 0.06$ | $p = 0.968$ | $X^2_1 = 16.89$ | $p = \mathbf{0.001}$ | $X^2_2 = 0.32$ | $p = 0.848$ |
| *RpSec27* | $X^2_1 = 2.37$ | $p = 0.123$ | $X^2_2 = 1.71$ | $p = 0.424$ | $X^2_1 = 0.02$ | $p = 0.879$ | $X^2_2 = 2.76$ | $p = 0.251$ |
| *RpGG* | $X^2_1 = 11.28$ | $p = \mathbf{<0.001}$ | $X^2_2 = 0.45$ | $p = 0.796$ | $X^2_1 = 3.38$ | $p = 0.065$ | $X^2_2 = 8.65$ | $p = \mathbf{0.013}$ |
| *RpMYR* | $X^2_1 = 4.69$ | $p = \mathbf{0.030}$ | $X^2_2 = 4.03$ | $p = 0.132$ | $X^2_1 = 6.50$ | $p = \mathbf{0.010}$ | $X^2_2 = 33.20$ | $p = \mathbf{<0.001}$ |
| *RpAD* | $X^2_1 = 7.51$ | $p = \mathbf{0.006}$ | $X^2_2 = 0.76$ | $p = 0.680$ | $X^2_1 = 4.10$ | $p = \mathbf{0.042}$ | $X^2_2 = 8.05$ | $p = \mathbf{0.017}$ |
| *RpMAPK4* | $X^2_1 = 3.42$ | $p = 0.064$ | $X^2_2 = 2.27$ | $p = 0.321$ | $X^2_1 = 0.50$ | $p = 0.476$ | $X^2_2 = 2.80$ | $p = 0.245$ |

**Fig. 4:**
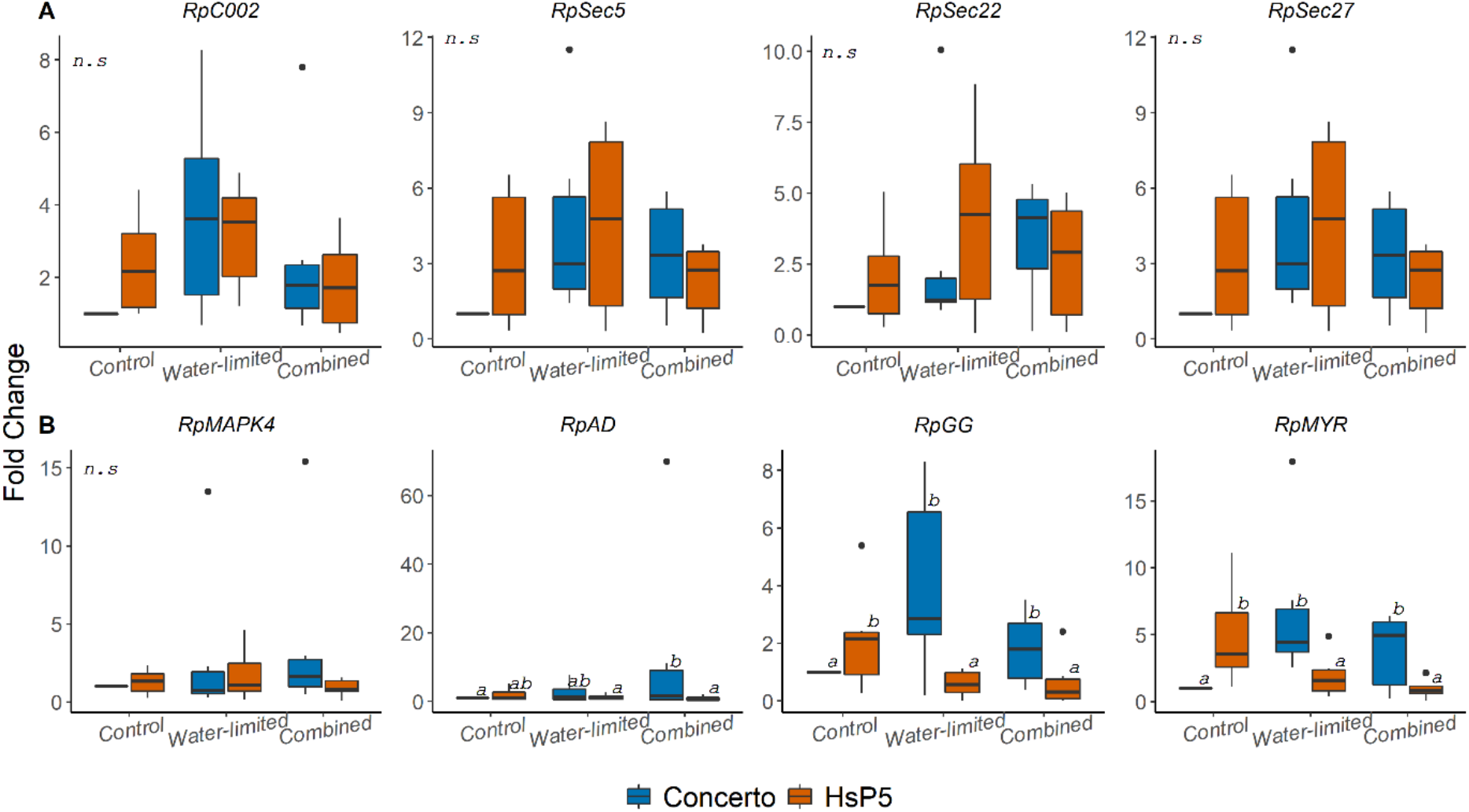
Expression patterns of *Rhopalosiphum padi* effector (A) and detoxification and stress-responsive (B) genes in adult aphids feeding on Concerto and Hsp5 under each water treatment. All gene expression values are relative to the mean expression of *RpCDC42*. Foldchange, 2-ΔΔCt, is relative to the expression of aphids feeding on Concerto under control conditions. Letters show significant differences based on least squares mean post-hoc analysis. *n* = 6.

However, the expression levels of the two detoxification genes, *RpGG* and *RpMYR*, and the stress-responsive gene *RpAD*, were differentially affected by the host plant type x water treatment interaction (Table 2; Fig. 4). Under control conditions, the two detoxification genes, *RpGG* and *RpMYR*, showed a two-fold higher expression levels in aphids feeding on Hsp5 compared with aphids feeding on Concerto (Fig. 4B). In response to drought stress, the expression levels of these two genes became elevated in aphids feeding on Concerto and reduced in aphids feeding on Hsp5 (Fig. 4B). Expression levels of the stress-responsive gene, *RpAD*, were elevated in aphids feeding on Concerto under the combined stress treatment compared with aphids feeding on Hsp5 under the same treatment (Fig. 4B).

## Discussion

Our results indicate that the physiological and molecular responses of plants to drought are, to a certain extent, mediated by constitutive resistance against sap-feeding insects, and that this has knock-on effects on the physiological and molecular responses of aphids to drought.

The observation that drought stress had a negative effect on plant physiological processes is in line with previous findings (Ivandic et al., 2000; González et al., 2008; Aslam et al., 2013; Zeppel et al., 2014) and the wider conclusion of meta-analyses (Cornelissen et al., 2008; Leybourne et al., 2021). We identified similar negative effects of drought stress on both plant types: the above- and below-ground dry mass decreased to similar levels when exposed to both drought stress conditions, although, the mass of Hsp5 was lower than Concerto under controlled conditions, indicating that Concerto suffered greater reduction due to drought stress than Hsp5. Root:shoot allometry increased in response to the drought treatments to a similar extent for both plant types. Differences were observed between the two plant types for root length and stomatal conductance, with Hsp5 possessing shorter roots and showing lower stomatal conductance than Concerto under all treatments examined. The physiological responses of both plant types to the drought stress treatments indicate that the level of drought stress experienced by the two plant types differed. The negative consequence of drought on the above- and below-ground mass of susceptible Concerto was greater than that observed for resistant Hsp5, indicating that the susceptible barley plants show a greater response to increasing drought stress than the resistant plants.

### Higher expression of the ABA-responsive gene, HvA1, in Hsp5 may contribute to drought-tolerance

The wild relatives of cereal crops have been widely reported to possess traits that confer abiotic tolerance and biotic resistance (Ellis et al., 2000; Zhao et al., 2010; Dempewolf et al., 2014; Al-Abdallat et al., 2017). Although both Hsp5 and Concerto showed reduced plant growth and impaired physiological processes under drought stress, there were some subtle differences between the responses of the two plant types. The stomatal conductance of Hsp5 was consistently lower than Concerto under control conditions and when exposed to drought stress. Control of stomatal pore size is a key mechanism through which plants can mediate transpiration and regulate water use efficiency during periods of low water availability (Brodribb and McAdam, 2011). This finding indicates that Hsp5 might harbour specific agronomic traits which could confer drought tolerance.

Abscisic acid (ABA)-mediated signalling contributes to drought tolerance (as reviewed by Sah et al., 2016). This phytohormone is primarily synthesised in the roots and plays a key role in mediating the responses of plants to drought stress, including controlling stomatal opening (Bandurska and Stroiński, 2003; Munemasa et al., 2015). *HvA1* encodes for an ABA-induced class 3 late embryogenesis-abundant (LEA) protein (Hong et al., 1992; Straub et al., 1994). In barley, *HvA1* has been linked to drought and salinity tolerance (Liang et al., 2012). Transcript levels of *HvA1* were decreased under the combined stress treatment compared with the water-limited treatment in both plant types. This likely reflects a slight difference in the water availability between the two drought stress treatments imposed, potentially due to increased contact with the soil in the combined stress treatment. Expression of the ABA-responsive gene, *HvA1* was consistently elevated in Hsp5 compared with Concerto, in line with our previous finding (Leybourne et al., 2019). This suggests that Hsp5 features elevated ABA signalling and indicates that this barley wild species exhibits enhanced drought tolerance. Indeed, previous transgenic studies have shown that transformation of rice and mulberry with *HvA1* results in a phenotype with increased tolerance of dehydration and drought (Chandra Babu et al., 2004; Lal et al., 2007; Checker et al., 2012). Additionally transgenic oat transformed with *HvA1* had enhanced salt tolerance (Oraby et al., 2005) and transgenic wheat exhibited enhanced water use efficiency under water stress (Sivamani et al., 2000). Barley wild relatives have also been widely reported to harbour drought tolerance traits (Lakew et al., 2011; Ahmed et al., 2013).

### Plant resistance against aphids mediates the extent to which aphid fitness is affected by drought stress

Drought stress also led to a reduction in aphid fitness (both the water-limited and combined stress treatments), compared with the control treatment, although the magnitude of reduction differed between Hsp5 and Concerto. Although not statistically significant, we identified a general trend that aphid fitness on Hsp5 was reduced to a lesser extent by drought treatment than fitness on Concerto, suggesting that resistance against aphids could influence the extent to which aphid fitness is negatively affected by plant drought stress. This would need to be examined in greater detail.

Gene transcript levels of *HvTHIO1* were increased ten-fold in Concerto under the two drought stress treatments compared with the control, whereas expression levels were consistent in Hsp5 under all treatments; providing evidence that the biochemical and molecular responses of the two plant types to drought stress differs. Similarly, drought stress increased thionin gene transcripts by two-fold in the barley cultivar Tobak (Ozturk et al., 2002). Higher expression of thionins has been associated with increased resistance against aphids (Delp et al., 2009; Mehrabi et al., 2014; Escudero-Martinez et al., 2017). Escudero-Martinez et al., (2017) showed that transient overexpression of *H. vulgare* thionin genes in *Nicotiana benthamiana* reduced the performance of the peach-potato aphid, *Myzus persicae*, indicating that thionins likely play an important role in plant defence against aphids. Elevated *HvTHIO1* levels in Concerto under the drought stress conditions could contribute towards the observed trend of greater reduction in aphid fitness (compared with Hsp5) under the water stress treatments, or could contribute to the increased expression of aphid detoxification genes when feeding on this plant type. Furthermore, a recent meta-analysis indicated that the defensive processes of plants are generally heightened under drought stress conditions and growth and vigour are reduced, with negative consequences for aphids (Leybourne et al., 2021); the findings of this research are in line with the conclusions drawn from this meta-analysis.

Although we observed an overall trend of lower aphid fitness on Hsp5 compared with Concerto, we did not detect any statistically significant differences in aphid fitness between the susceptible variety of barley and partially-resistant Hsp5, as has been reported in the literature (Leybourne et al., 2019; Delp et al., 2009). One potential explanation for this could be the difference in aphid measurements taken between this study and previous studies (e.g., Leybourne et al., 2019). For example, previous studies used detailed life-history measurements to characterise aphid fitness on Hsp5 and a susceptible comparison, whereas in this study, due to the experimental setup of the drought experiment, we were constrained to generic measures of insect abundance over a defined time-period.

### Induction of aphid detoxification genes under drought stress is influenced by host plant resistance

The four aphid effector genes examined were not significantly affected by drought stress or host plant resistance. Similarly, exposure to different feeding environments, such as artificial diet, host and non-host plants does not affect the expression or *R. padi* effectors (Escudero-Martinez et al., 2020). Although we did not observe changes in aphid effector gene expression, it is possible that secretion or function of effectors is affected by drought stress or exposure to resistant plants, affecting their role in promoting aphid virulence.

The two detoxification genes examined (*RpGG* and *RpMYR*, encoding λ-glutamylcysteine synthetase and a myrosinase-like gene, respectively) were two- and five-fold more highly expressed, respectively, in aphids feeding on Hsp5 than Concerto under control conditions. In the soybean aphid, *Aphis glycine* Matsumara, expression of genes associated with glutathione biosynthesis and the glutathione reduction pathway were similarly up-regulated in response to resistant soybean (Bansal et al., 2014). However, *RpGG* and *RpMYR* expression decreased in aphids on Hsp5 when exposed to the two drought stress treatments compared with the control, while expression increased in aphids on Concerto when exposed to the drought stress treatments. λ-glutamylcysteine synthetase (also known as glutamate cysteine ligase) is a key component of glutathione biosynthesis (Forman et al., 2009). Glutathiones are key antioxidants involved in the detoxification of xenobiotic compounds (Forman et al., 2009) and are found in many plant, animal, and fungi species. In aphids, these are routinely involved in the detoxification of plant secondary metabolites and insecticides, in a process which involves glutathione-S transferase (Francis et al., 2005; Jeschke et al., 2016; Balakrishnan et al., 2018). Similarly, myrosinase is also involved in the detoxification of plant defensive compounds (Francis et al., 2002).

As cereals do not produce glucosinolates, the role of myrosinase genes in cereal aphids is unknown. These genes are highly expressed in various cereal and grass aphid species (Escudero-Martinez et al., 2017; Koch et al., 2019), and are predicted to play an active role in facilitating successful aphid infestation through the inactivation of ingested plant defensive compounds. The increased expression of *RpGG* and *RpMYR* in *R. padi* feeding on Concerto under drought stress conditions parallels the observed increase in *HvTHIO1* expression in Concerto under these conditions, suggesting that *RpGG* and *RpMYR* transcripts may have a function in aphid responses to plant resistance and detoxification of plant defensive compounds, as suggested for other aphid species (Francis et al., 2002; Bansal et al., 2014).

The EPG data indicated that aphid responses to drought stress are independent of host plant resistance. Under drought, the ingestion of phloem sap was restricted and xylem ingestion was promoted for aphids feeding on both plant types. A biochemical response of plants to drought is an increase in the solute concentration of the phloem (Xiong and Zhu, 2002; Sevanto, 2014), which can result in increased phloem osmolarity, with detrimental effects on aphid fitness. Decreases in concentrations of essential amino acids in the phloem have also been reported (Lin et al., 2021). In response to increased phloem osmolarity it has been hypothesised that aphids mix xylem and phloem as a form of osmoregulation (Pompon et al., 2011), with increased periods of xylem ingestion acting to reduce the negative effects of high phloem osmolarity. Therefore, even though the molecular and biochemical responses of two plant types differ in response to drought stress, the effect of these changes on aphid feeding behaviour is similar, although, the impact on the quantity of phloem ingested (and therefore nutritional value) was not measured in the present study.

## Conclusion

Here we examined how plant resistance traits, exposure to drought, and infestation with aphids interact and how this impacts the physiological and molecular responses of plants. We focussed on a well characterised plant-aphid system comprising a susceptible barley variety (Concerto) and a wild progenitor of barley with partial-resistance against aphids (Hsp5). The molecular responses of the aphids suggested that responses to drought stress may be mediated by the two contrasting plant types, with the expression of detoxification genes increasing on susceptible plants under drought, but decreasing when aphids were feeding on aphid-resistant plants. We also characterised some of the molecular determinants that could have contributed to this observation at the plant-level and found that this is likely associated with an increase in the expression of defence-related genes in susceptible barley under drought stress conditions, particularly the thionin gene *HvTHIO1*. This study provides new insight into the complex interactions between plants, aphids, and the environment and shows that plant resistance traits are important factors to consider when examining plant-insect interactions under environmental stress.

## Acknowledgements

DJL was funded by the James Hutton Institute and the Universities of Aberdeen and Dundee through a Scottish Food Security Alliance (Crops) PhD studentship. AJK, KB, AT, JIBB, and TAV were supported by the strategic research programme funded by the Scottish Government’s Rural and Environment Science and Analytical Services Division. The authors would like to thank Dr Jenny Slater (The James Hutton Institute) for helping with data collection in the drought experiments and Dr Katharine Preedy (Biomathematics and Statistics Scotland) for providing statistical advice.

## Author Contributions

JIBB, AJK, TAV, and DJL conceived and designed the experiments. DJL performed the experiments. DJL, KB, and AT performed the tension table monitoring experiments needed to construct the calibration curve. JIBB, AJK, TAV, and DJL analysed the data. JIBB and DJL wrote the manuscript with input from AJK, and TAV. All authors read and approved the final manuscript.

