## Supplementary file and tables for "Drought stress increases the expression of barley defence genes with negative consequences for infesting cereal aphids"

***Supplementary File 1:***

***Materials and methods for the development of the drought stress treatments***

*Soil physical properties of growth medium*

Soil was excavated from the Mid-Pilmore field at the James Hutton Institute, Dundee, UK in 2015. Soil physical characteristics were assessed to obtain water release curves and soil strength at specific matric potentials (Valentine et al 2012). Briefly, excavated soil was dried and sieved < 2mm to remove large stones and debris. A subsample of sieved soil (*n = 6*) was compacted into cylindrical cores (5 cm diameter, 5 cm height, internal volume of 95.03 $cm^3$) at a compaction density of 1.15 g $cm^3$ or 1.25 g $cm^3$ dry bulk density and saturated to field capacity. The matric potential (MP) of the soil in these cores was adjusted to -5 kPa using silica sand tables and decreased to -300 kPa on ceramic suction plates, and then a subsample from each core was taken through to -1500 kPa on pressure plates (ELE Ltd, UK). In this way, the MP was adjusted serially through 0, -1, -5, -20, -50, -300, and -1500 kPa (Richards and Weaver, 1944). Soil strength was determined by taking penetrometer (Instron model 5544; Instron, MA, USA.) recordings at -5, -20, and -300 kPa stages of the water release curve using a needle penetrometer (1 mm diameter, $30^o$ cone angle, 4 mm $min^{-1}$ penetration rate; averaged at 0.75 mm intervals from 4.5-9.75 mm depth range).

Three water treatments were selected based on the calibrated gravimetric moisture content (gMC) of these soil cores. The control treatment was designed to be below field capacity (*c.* 40 % field capacity) with an acceptable level of aeration and a compaction density and soil strength which would not impede root growth (1.15 g $cm^3$ soil density; *c.* 1.06 MPa soil strength), and two drought stress treatments were designed so that the gMC was lower than the control treatment but above the permanent wilting point (*c.* 30 % field capacity): one treatment (water-limited treatment) had a compaction density similar to the Control treatment (1.15 g $cm^3$ soil density; 2.24 MPa soil strength); and a second treatment (combined treatment) had an increased soil compaction density (1.25 g $cm^3$ soil density; 3.92 MPa soil strength) which aimed to impede root growth. Soil was mixed to a gMC of *c.* 0.22 g $cm^3$ for the control treatment (estimated MP = 15-20 kPa) and *c.* 0.16 g $cm^3$ for the two drought stress treatments (estimated MP for the water-limited treatment = 100-350 kPa, estimated MP for the combined treatment = 400-500 kPa). The soil MP was recorded routinely using a tensiometer (model SW-T5, Delta-T Devices, UK) to ensure the soil matrix remained around the desired levels, plants were watered when required to maintain this level.

*Validation of devised water treatments*

To test the effect of the devised water stress treatments a preliminary study was undertaken. Soil was compacted into cylindrical Perspex tubes (internal volume *c.* 950.03 $cm^3$; *c.* 5.00 cm diameter) at a density of 1.15 g $cm^3$ or 1.25 g $cm^3$ (depending on treatment) and Concerto and Hsp5 plants were grown under control, water-limited, or combined conditions ($n = 3$). Stomatal conductance (mmol $cm^2$ $s^{-1}$) was measured at three time-points during plant development and plants were destructively harvested and above-ground dry mass was recorded when they reached the three-leaf stage of development (Zadoks et al., 1974).

*Statistical analysis of soil physical data*

gMC, volumetric water content (VWC), air filled porosity (AFP), soil strength (determined by measuring the penetrometer resistance, PR, using an Instron penetrometer), and the volume of soil occupied by air or pores were analysed in separate Type II ANOVA models in response to the MP, soil compaction, and the interaction. Differences of least squares means with Tukey correction was used as a post-hoc test on each model. Fitted-residual plots of the final models were assessed for model suitability.

*Statistical analysis of preliminary experiment: water treatment validation*

Stomatal aperture was analysed as a response variable against plant type, water treatment, and the interaction in a mixed effects model fitted with a temporal correlation structure for repeated measures analysis. Plant shoot dry mass was analysed by Type II ANOVA in response to plant type, water treatment, and the interaction. Fitted-residual plots of the final models were assessed for model suitability.

### ***Results of water treatment validation***

*Soil physical analysis results*

To devise suitable water treatments, excavated field soil was taken through a series of MPs to generate a water release curve to provide information on the physical characteristics of the soil and to elucidate the water retention properties of the soil matrix. The gMC of soil in the cores was significantly affected by the matric suction applied ($F_{6,70}$ = 1048.68; p = <0.001), the level of soil compaction ($F_{1,70}$= 5.17; p = 0.025) and the applied suction x soil compaction interaction ($F_{6,70}$= 10.01; p = <0.001), with the gMC of the soil matrix decreasing steadily as the MP became more negative (Fig. SF1-1A). Similarly, the VWC of the soil cores was significantly affected by the suction applied ($F_{6,70}$ = 1133.75; p = <0.001), soil compaction ($F_{1,70}$= 10.74; p = 0.001) and the applied suction x soil compaction interaction ($F_{6,70}$= 5.97; p = <0.001), with the volume of water retained in the soil matrix decreasing as MP became more negative (increasing suction) (Fig. SF1-1B). To examine how soil compaction and altered MP might affect plant root growth, the soil strength (PR) was determined. Soil strength significantly increased alongside more negative MPs ($F_{2,30}$ = 44.79; p =<0.001) and soil compaction ($F_{1,30}$ = 21.84; p = <0.001; Fig. SF1-1C), with soil strength around two-fold higher in the more dense soil matrix (1.25 g $cm^3$) compared with the less dense soil (1.15 g $cm^3$) at the same MP (Fig. SF1-1C). The volume of the soil matrix which was occupied by different size categories of pores, as estimated by the difference in VWC at different MPs, showed different volumes of the different pore size categories ($F_{5,60}$ = 126.81; p = <0.001), different overall volume of pores between the soil compaction treatments ($F_{1,60}$ = 4.47; p = 0.038) and a significant interaction ($F_{5,60}$ = 7.62; p = <0.001; Fig. SF1-1D), such that the compacted soil had significantly fewer larger pores than the low compaction soil cores.

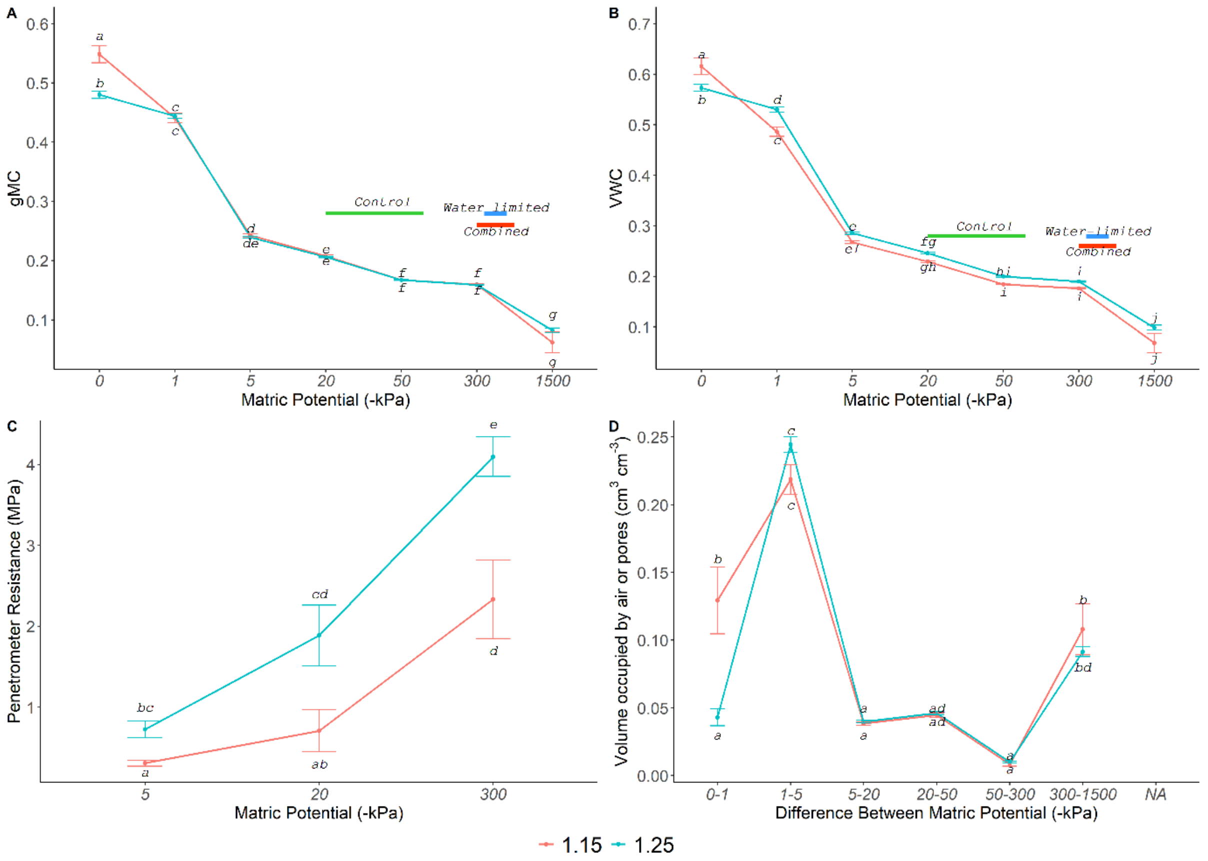


***Fig. SF1-1:*** *Tension table results from the soil cores used to devise the water treatments. High compaction cores (blue), low compaction cores (red). Graphs show soil physical characteristics* n = 6. *A) gMC, B) VWC, C) PR, D) the volume of soil occupied by pores of different size categories. Error bars represent mean ± se. Horizontal lines displayed on panels A and B show the upper and lower interquartile range of the MP (-kPa) of the soil matrices used throughout the subsequent water limitation experiments for the three devised treatments: Control, Water- limited (Water), and Combined.*

*The devised drought stress treatments successfully reduce plant growth and decrease stomatal aperture*

A preliminary investigation was carried out in order to validate the devised water treatments. Shoot dry mass and stomatal conductance of Concerto and Hsp5 were measured when growing under the control, water-limited, or combined stress treatments. Shoot dry mass was significantly reduced under the imposed drought stress treatments ($F_{2,17}$ = 8.05; p = 0.003; Table SF1-1; Table SF1-2), with no difference detected between plant types ($F_{2,17}$ = 2.09; p = 0.153) and there was no interaction ($F_{4,17}$ = 1.15; p = 0.366). Tukey post-hoc analysis indicated that plant mass differed between the control and water-limited treatment (p = 0.0.013) and the control and combined treatment (p = 0.046), but not between the water-limited treatment and the combined treatment (p = 0.875). The applied drought stress also had a negative effect on stomatal conductance (Table SF1-1; Table SF1-2). Stomatal conductance was reduced due to water stress treatment at time-point one ($F_{2,17}$ = 9.09; p = 0.002), two ($F_{2,17}$ = 6.85; p = 0.006), and three ($F_{2,17}$ = 9.59; p = 0.001). Post-hoc analysis indicated that for the first time-point this difference was between the control and the water-limited treatment (p = 0.003), for time-point two this was between the control and combined treatment (p = 0.015), and for the third time-point this was between the control and the water-limited (p = 0.040) and control and the combined treatments (p = 0.009).

***Table SF1-1:*** *Results of preliminary water-stress trial (*n = 3*)*

| Measurement | Mean Value | | | | | |
|---|---|---|---|---|---|---|
| | Concerto Control | Hsp5 Control | Concerto Water-limited | Hsp5 Water-limited | Concerto Combined | Hsp5 Combined |
| Shoot dry mass (mg) | 109.00 | 125.66 | 74.00 | 77.33 | 97.66 | 59.00 |
| Stomatal Conductance (mmol cm$^2$ s$^{-1}$) Measurement 1 | 88.00 | 63.50 | 28.30 | 21.70 | 48.00 | 58.50 |
| Stomatal Conductance Measurement 2 | 107.30 | 78.50 | 60.30 | 33.90 | 15.90 | 26.50 |
| Stomatal Conductance Measurement 3 | 131.30 | 88.30 | 84.30 | 46.70 | 45.40 | 46.00 |

***Table SF1-2:*** *Statistical results of preliminary water-stress trial (*n = 3*). Bold text indicates significant p values.*

| Measurement | Statistical Results: $F_{(df)}$; p-value | | |
|---|---|---|---|
| | Plant Type | Water Treatment | Plant Type x Water Treatment |
| Shoot dry mass (mg) | $F_{2,17}$ = 2.09; p = 0.153 | $F_{2,17}$ = 8.05; p = 0.003 | $F_{4,17}$ = 1.15; p = 0.366 |
| Stomatal Conductance (mmol cm$^2$ s$^{-1}$) Measurement 1 | $F_{2,17}$ = 0.71; p = 0.506 | $F_{2,17}$ = 9.09; **p = 0.002** | $F_{4,17}$ = 2.83; p = 0.056 |
| Stomatal Conductance Measurement 2 | $F_{2,17}$ = 1.44; p = 0.264 | $F_{2,17}$ = 6.85; **p = 0.006** | $F_{4,17}$ = 2.76; p = 0.061 |
| Stomatal Conductance Measurement 3 | $F_{2,17}$ = 4.79p = **0.022** | $F_{2,17}$ = 9.59; **p = 0.001** | $F_{4,17}$ = 2.26; p = 0.104 |

### *Supplementary Tables: S1-S7*

**Table S1:** Barley RT-qPCR primers used in this study

| Gene | Name | Accession no. | Forward primer | Reverse Primer | Efficiency | Statistical Transformation | Reference |
|---|---|---|---|---|---|---|---|
| *HvGR* | glycine-rich protein RNA-binding protein | TC146685 | cgcccagttatc catccatcta | aaaaacaccaca ggaccggac | 93% | N/A – reference gene | Faccioli et al. 2007 |
| *HvEF-1-α* | Elongation factor 1 α | TC146566 | atgattcccacc aagcccat | acaccaacagcc acagtttgc | 87% | N/A – reference gene | Faccioli et al. 2007 |
| *HvLOX2* | Predicted Lipoxygenase | AK357253.1 | atgtcctatccc acgacacc | agtgcgtcctcag ccagt | 103% | Not transformed | Escudero-Martinez et al 2017 |
| *HvA1* | ABA-inducible late embryogenesis abundant protein | X13498.1 | atgggagggg acaacacc | ggaaattaagcg cgaacg | 91% | Log transformed | Designed for this study |
| *HvNPR1* | Non-expresser of pathogenesis-related genes 1-Like | MLOC_6492 2.1 | ttgataacatct agaggcaatg ct | tgcgtgaaactgtt cgagag | 84% | Square-root transformed | Designed for this study |
| *HvERF1* | Ethylene-response factor 1 | HQ328941.1 | ctatataatgatt gggtgcatgttg | ggcatatgaccca aggtgtt | 87% | Square-root transformed | Designed for this study |
| *HvTHIO 1* | Thionin 1 | AK359149 | tatggccaagg tcgttttgt | cataactaagatg atacatttgcttcg | 118% | Log transformed | Escudero-Martinez et al 2017 |

**Table S2:** Aphid RT-qPCR primers used in this study.

| Gene | Name | Forward primer | Reverse Primer | Efficiency | Statistical Transformation | Reference |
|---|---|---|---|---|---|---|
| *RpCDC42* | Cell Division Control Protein 42 | ttttggttgatttagacggatgt | agttgatggatgaagaattactggt | 90% | N/A – reference gene | Designed for this study |
| *RpGG* | λ-glutamylcysteine synthetase | cctggactaataaatttttacggaat | ggccgttttccaggttttat | 87% | Log transformed | Designed for this study |
| *RpAD* | Alcohol Dehydrogenase | accaatcactacacatttaacaggtc | tgaagacaatgaagatgggcta | 90% | Log transformed | Designed for this study |
| *RpMYR* | Myrosinase-like | gatttgacgcatcacaggaa | ggcaaatgattgctgttgtc | 86% | Log transformed | Designed for this study |
| *RpMAPK4* | Mitogen Activated Protein Kinase 4 | tcctgttctcgtgccattct | acaaaaacgtaaagatcaattggaa | 70% | Log transformed | Designed for this study |
| *RpCOO2* | Effector COO2 | cgctcactactcgatgggtat | cgtgacgtctacctctctcaaa | 106% | Log transformed | Designed for this study |
| *RpSec5* | Secreted Effector 5 | caggaaaagttgactattctgctgta | gagccatttgcttttagacttga | 115% | Log transformed | Designed for this study |
| *RpSec22* | Secreted Effector 22 | tgctcaaggtgcatacgact | agtgttttaaagctttttcgaatctt | 114% | Log transformed | Designed for this study |
| *RpSec27-1* | Secreted Effector 27-1 | atatgacatggaaattcaaagacg | cttcttcatcaagtaattgacgttct | 80% | Log transformed | Designed for this study |

**Table S3:** Mean values ± standard error of the mean for plant physiological parameters measured throughout the drought stress experiment

| **Variable measured** | **Control** | | | | **Water-limited** | | | | **Compounding** | | | |
|---|---|---|---|---|---|---|---|---|---|---|---|---|
| | **Concerto** | | **Hsp5** | | **Concerto** | | **Hsp5** | | **Concerto** | | **Hsp5** | |
| | **Aphid infested** | **Uninfested** | **Aphid infested** | **Uninfested** | **Aphid infested** | **Uninfested** | **Aphid infested** | **Uninfested** | **Aphid infested** | **Uninfested** | **Aphid infested** | **Uninfested** |
| Development time to true-leaf stage (days) ¶ | 11.36 ± 0.25 | 11.77 ± 0.38 | 12.86 ± 0.55 | 12.85 ± 0.66 | 17.14 ± 1.37 | 17.57 ± 1.23 | 17.00 ± 1.37 | 16.85 ± 1.48 | 16.00 ± 1.35 | 17.15 ± 1.39 | 17.93 ± 1.34 | 18.67 ± 1.69 |
| Mean stomatal conductance (mmol $s^{-1}$) ¶ | 182.56 ± 15.93 | 188.60 ± 12.88 | 128.20 ± 9.53 | 147.10 ± 9.78 | 145.10 ± 19.74 | 102.95 ± 9.81 | 91.09 ± 11.34 | 71.15 ± 5.89 | 121.00 ± 13.30 | 171.60 ± 13.14 | 79.42 ± 6.43 | 105.76 ± 17.21 |
| Above ground dry mass (mg) | 237.20 ± 1.86 | 202.10 ± 15.96 | 156.60 ± 11.55 | 168.20 ± 13.44 | 62.06 ± 7,79 | 56.53 ± 17.39 | 54.09 ± 11.32 | 61.42 ± 10.66 | 69.16 ± 12.95 | 61.68 ± 12.72 | 42.72 ± 9.16 | 40.22 ± 14.30 |
| Below ground dry mass (mg) | 94.90 ± 13.24 | 85.61 ± 14.18 | 78.72 ± 8.90 | 75.28 ± 8.07 | 41.84 ± 4.51 | 42.42 ± 6.01 | 47.39 ± 8.48 | 42.44 ± 6.02 | 45.27 ± 5.06 | 34.53 ± 3.59 | 33.15 ± 4.22 | 33.37 ± 6.19 |
| Root length (mm) | 518.30 ± 15.10 | 480.90 ± 36.91 | 486.10 ± 6.89 | 483.10 ± 9.75 | 427.60 ± 14.98 | 414.10 ± 10.94 | 378.20 ± 15.31 | 374.80 ± 28.57 | 407.90 ± 21.81 | 343.10 ± 36.22 | 318.60 ± 24.22 | 333.00 ± 17.52 |
| Root:Shoot Allometry | 0.39 ± 0.04 | 0.39 ± 0.06 | 0.52 ± 0.05 | 0.45 ± 0.03 | 0.79 ± 0.10 | 1.52 ± 0.47 | 1.43 ± 0.37 | 1.12 ± 0.34 | 0.94 ± 0.17 | 1.03 ± 0.23 | 1.01 ± 0.14 | 1.46 ± 0.29 |

¶ These parameters were taken before aphid infestation

**Table S4:** Mean values ± standard error of the mean for aphid fitness parameters measured throughout the drought stress experiment

| Variable measured | Control | | Water-limited | | Combined | |
|---|---|---|---|---|---|---|
| | Concerto | Hsp5 | Concerto | Hsp5 | Concerto | Hsp5 |
| Cumulative fecundity | 97.07 ± 8.82 a | 89.15 ± 7.18 ab | 63.07 ± 8.27 d | 65.21 ± 6.20 bc | 62.62 ± 10.35 d | 65.62 ± 10.89 cd |
| Freeze-dried mass of cumulative progeny (mg) | 1.99 ± 0.27 a | 1.63 ± 0.22 a | 1.07 ± 0.23 b | 1.25 ± 0.21 b | 1.20 ± 0.37 b | 0.96 ± 0.22 b |

Letters indicate significant differences based on least squares mean post-hoc analysis.

**Table S5:** Mean values ± standard error of the mean for aphid feeding parameters measured throughout the EPG drought stress experiment. Parameter descriptions are provided in the table footer; full parameter details and their calculation is available in the macro developed by Schliephake et al., 2013.

| Parameter Measured | Control | | | | | | Water-limited | | | | | | Combined | | | | | |
|---|---|---|---|---|---|---|---|---|---|---|---|---|---|---|---|---|---|---|
| | ***Concerto*** | | | ***Hsp5*** | | | ***Concerto*** | | | ***Hsp5*** | | | ***Concerto*** | | | ***Hsp5*** | | |
| | *Mean* | *SE* | *n* | *Mean* | *SE* | *n* | *Mean* | *SE* | *n* | *Mean* | *SE* | *n* | *Mean* | *SE* | *n* | *Mean* | *SE* | *n* |
| n_Np | 4.78 | 0.81 | 9 | 5.90 | 1.66 | 10 | 8.14 | 2.11 | 7 | 6.14 | 1.98 | 7 | 3.00 | 1.21 | 8 | 6.18 | 1.38 | 11 |
| a_Np (s) | 440.24 | 175.10 | 9 | 517.85 | 260.00 | 10 | 390.46 | 77.19 | 7 | 611.31 | 188.81 | 7 | 634.97 | 178.16 | 8 | 253.66 | 58.99 | 11 |
| m_Np (s) | 497.14 | 244.01 | 9 | 502.11 | 259.90 | 10 | 218.15 | 41.67 | 7 | 467.93 | 221.75 | 7 | 566.63 | 176.64 | 8 | 172.42 | 48.37 | 11 |
| s_Np (s) | 1445.70 | 485.00 | 9 | 2321.02 | 716.52 | 10 | 3432.23 | 1190.85 | 7 | 3907.34 | 1389.47 | 7 | 1378.50 | 454.25 | 8 | 1772.59 | 543.56 | 11 |
| n_Pr | 4.78 | 0.81 | 9 | 5.90 | 1.66 | 10 | 8.14 | 2.11 | 7 | 6.14 | 1.98 | 7 | 3.00 | 1.21 | 8 | 6.18 | 1.38 | 11 |
| n_bPr | 1.33 | 0.58 | 9 | 2.00 | 0.88 | 10 | 2.86 | 1.37 | 7 | 2.14 | 1.34 | 7 | 0.50 | 0.27 | 8 | 1.73 | 0.52 | 11 |
| a_Pr (s) | 6425.48 | 1980.49 | 9 | 6911.31 | 1959.55 | 10 | 3415.26 | 856.17 | 7 | 6075.91 | 2657.11 | 7 | 13335.49 | 2930.24 | 8 | 7279.06 | 2291.89 | 11 |
| m_Pr (s) | 4809.95 | 2268.63 | 9 | 6200.56 | 2121.15 | 10 | 1879.22 | 744.66 | 7 | 4951.09 | 2853.62 | 7 | 12168.43 | 3423.38 | 8 | 5664.88 | 2521.64 | 11 |
| s_Pr (s) | 20154.30 | 485.00 | 9 | 19278.98 | 716.52 | 10 | 18167.77 | 1190.85 | 7 | 17692.66 | 1389.47 | 7 | 20221.50 | 454.25 | 8 | 19827.41 | 543.56 | 11 |
| n_C | 9.44 | 1.46 | 9 | 14.30 | 3.12 | 10 | 13.86 | 2.05 | 7 | 11.00 | 2.18 | 7 | 8.38 | 2.60 | 8 | 10.00 | 1.89 | 11 |
| a_C (s) | 708.30 | 121.41 | 9 | 658.31 | 76.21 | 10 | 555.13 | 85.88 | 7 | 653.34 | 85.11 | 7 | 707.89 | 194.50 | 8 | 992.26 | 243.01 | 11 |
| m_C (s) | 494.45 | 99.06 | 9 | 431.15 | 65.83 | 10 | 384.77 | 107.65 | 7 | 457.75 | 95.35 | 7 | 355.14 | 104.66 | 8 | 726.12 | 271.50 | 11 |
| s_C (s) | 6754.46 | 1501.65 | 9 | 8282.82 | 1288.10 | 10 | 7124.32 | 931.45 | 7 | 6690.09 | 1298.18 | 7 | 5018.19 | 1635.23 | 8 | 7368.96 | 1341.27 | 11 |
| n_F | 0.11 | 0.11 | 9 | 0.60 | 0.31 | 10 | 0.43 | 0.30 | 7 | 0.00 | 0.00 | 7 | 0.00 | 0.00 | 8 | 0.09 | 0.09 | 11 |
| a_F (s) | 1329.90 | NA | 1 | 1391.14 | 609.89 | 4 | 1737.05 | 1522.03 | 2 | NA | NA | 0 | NA | NA | 0 | 970.00 | NA | 1 |
| m_F (s) | 1329.90 | NA | 1 | 1360.19 | 623.61 | 4 | 1737.05 | 1522.03 | 2 | NA | NA | 0 | NA | NA | 0 | 970.00 | NA | 1 |
| s_F (s) | 147.77 | 147.77 | 9 | 679.89 | 343.66 | 10 | 961.88 | 926.54 | 7 | 0.00 | 0.00 | 7 | 0.00 | 0.00 | 8 | 88.18 | 88.18 | 11 |
| n_G | 1.22 | 0.36 | 9 | 2.00 | 0.49 | 10 | 2.86 | 0.80 | 7 | 2.43 | 0.53 | 7 | 2.75 | 0.70 | 8 | 2.45 | 0.43 | 11 |
| a_G (s) | 4216.80 | 1090.90 | 6 | 3232.47 | 1474.98 | 9 | 5180.55 | 1297.58 | 7 | 6570.16 | 2427.02 | 7 | 8263.45 | 2643.65 | 8 | 6921.19 | 2013.45 | 11 |
| m_G (s) | 4066.33 | 1162.48 | 6 | 3114.16 | 1484.43 | 9 | 4454.13 | 1565.98 | 7 | 6611.69 | 2420.80 | 7 | 7830.34 | 2778.77 | 8 | 6649.78 | 2068.88 | 11 |
| s_G (s) | 4471.89 | 1390.64 | 9 | 4213.96 | 1380.69 | 10 | 9813.03 | 1864.54 | 7 | 10814.22 | 2053.22 | 7 | 13829.24 | 2334.24 | 8 | 11728.10 | 1867.11 | 11 |
| t_1G (s) | 12648.64 | 2402.21 | 9 | 7076.28 | 1943.23 | 10 | 4005.69 | 1483.92 | 7 | 3624.48 | 1219.78 | 7 | 3286.27 | 1031.63 | 8 | 5973.67 | 1503.09 | 11 |
| nPr_1G | 1.44 | 0.44 | 9 | 3.40 | 1.17 | 10 | 2.14 | 0.51 | 7 | 2.00 | 0.44 | 7 | 1.50 | 0.27 | 8 | 4.09 | 1.12 | 11 |
| n_sgE1 | 1.56 | 0.47 | 9 | 3.20 | 0.90 | 10 | 2.00 | 0.65 | 7 | 1.71 | 0.68 | 7 | 1.88 | 0.81 | 8 | 0.82 | 0.33 | 11 |
| a_sgE1 (s) | 140.09 | 51.75 | 6 | 61.65 | 15.20 | 8 | 24.98 | 1.39 | 5 | 37.75 | 11.62 | 5 | 51.13 | 20.59 | 5 | 59.79 | 21.63 | 5 |
| m_sgE1 (s) | 138.25 | 52.27 | 6 | 57.17 | 14.72 | 8 | 23.89 | 1.71 | 5 | 36.91 | 11.94 | 5 | 49.60 | 20.95 | 5 | 60.27 | 21.51 | 5 |
| s_sgE1 (s) | 200.16 | 83.08 | 9 | 192.18 | 57.52 | 10 | 51.22 | 18.19 | 7 | 72.46 | 36.92 | 7 | 85.60 | 38.21 | 8 | 42.59 | 17.69 | 11 |
| mx_sgE1 (s) | 222.14 | 96.96 | 6 | 107.07 | 24.19 | 8 | 36.23 | 3.84 | 5 | 53.34 | 21.84 | 5 | 77.47 | 30.11 | 5 | 66.27 | 20.64 | 5 |
| n_frE1 | 2.33 | 0.44 | 9 | 2.90 | 0.78 | 10 | 0.71 | 0.29 | 7 | 1.00 | 0.31 | 7 | 1.25 | 0.41 | 8 | 0.73 | 0.24 | 11 |
| a_frE1 (s) | 240.40 | 59.43 | 9 | 106.41 | 24.08 | 8 | 63.55 | 7.73 | 4 | 37.24 | 15.01 | 5 | 485.82 | 426.31 | 5 | 146.32 | 54.01 | 6 |
| m_frE1 (s) | 240.93 | 59.48 | 9 | 103.37 | 24.94 | 8 | 63.55 | 7.73 | 4 | 37.24 | 15.01 | 5 | 485.47 | 426.40 | 5 | 146.32 | 54.01 | 6 |
| s_frE1 (s) | 410.07 | 53.47 | 9 | 242.53 | 55.29 | 10 | 43.67 | 16.54 | 7 | 43.37 | 25.10 | 7 | 340.80 | 265.51 | 8 | 107.90 | 47.64 | 11 |

| | | | | | | | | | | | | | | | | | | |
|---|---|---|---|---|---|---|---|---|---|---|---|---|---|---|---|---|---|---|
| mx_frE1 (s) | 287.14 | 60.29 | 9 | 145.35 | 28.87 | 8 | 64.13 | 7.45 | 4 | 39.51 | 15.16 | 5 | 508.52 | 420.69 | 5 | 158.81 | 56.11 | 6 |
| n_E1 | 3.89 | 0.68 | 9 | 6.10 | 1.42 | 10 | 2.71 | 0.71 | 7 | 2.71 | 0.78 | 7 | 3.13 | 1.11 | 8 | 1.55 | 0.49 | 11 |
| a_E1 (s) | 192.53 | 44.06 | 9 | 83.05 | 20.20 | 9 | 34.88 | 4.59 | 6 | 42.14 | 10.26 | 7 | 198.05 | 154.73 | 5 | 125.22 | 46.78 | 7 |
| m_E1 (s) | 174.50 | 41.95 | 9 | 75.58 | 19.35 | 9 | 33.02 | 4.72 | 6 | 37.81 | 10.08 | 7 | 66.67 | 31.32 | 5 | 121.99 | 46.80 | 7 |
| s_E1 (s) | 610.23 | 115.92 | 9 | 434.72 | 98.20 | 10 | 94.90 | 25.60 | 7 | 115.83 | 41.09 | 7 | 426.40 | 293.40 | 8 | 150.50 | 54.75 | 11 |
| mx_E1 (s) | 317.54 | 70.78 | 9 | 149.45 | 30.68 | 9 | 52.20 | 8.90 | 6 | 57.34 | 16.57 | 7 | 508.52 | 420.69 | 5 | 156.24 | 47.49 | 7 |
| n_E12 | 2.22 | 0.43 | 9 | 2.80 | 0.76 | 10 | 0.57 | 0.20 | 7 | 1.00 | 0.31 | 7 | 1.13 | 0.40 | 8 | 0.73 | 0.24 | 11 |
| a_E12 (s) | 5614.29 | 1732.52 | 9 | 4017.27 | 2257.03 | 8 | 380.29 | 276.41 | 4 | 94.54 | 41.40 | 5 | 1248.26 | 539.54 | 5 | 920.79 | 637.17 | 6 |
| m_E12 (s) | 5176.86 | 1834.77 | 9 | 3659.61 | 2324.70 | 8 | 380.29 | 276.41 | 4 | 94.54 | 41.40 | 5 | 1142.39 | 572.02 | 5 | 920.79 | 637.17 | 6 |
| s_E12 (s) | 8580.01 | 1688.47 | 9 | 5910.13 | 1965.48 | 10 | 217.31 | 166.53 | 7 | 115.89 | 69.37 | 7 | 1288.47 | 633.16 | 8 | 599.57 | 370.18 | 11 |
| mx_E12 (s) | 7568.16 | 1686.53 | 9 | 5912.05 | 2172.15 | 8 | 380.29 | 276.41 | 4 | 103.05 | 45.37 | 5 | 1940.79 | 843.51 | 5 | 983.06 | 633.05 | 6 |
| n_E2 | 2.22 | 0.43 | 9 | 2.80 | 0.76 | 10 | 0.57 | 0.20 | 7 | 1.00 | 0.31 | 7 | 1.13 | 0.40 | 8 | 0.73 | 0.24 | 11 |
| a_E2 (s) | 5364.77 | 1677.65 | 9 | 3908.29 | 2239.00 | 8 | 303.86 | 274.17 | 4 | 57.30 | 27.23 | 5 | 752.40 | 407.80 | 5 | 774.47 | 586.56 | 6 |
| m_E2 (s) | 4924.84 | 1780.37 | 9 | 3545.10 | 2307.48 | 8 | 303.86 | 274.17 | 4 | 57.30 | 27.23 | 5 | 655.40 | 421.11 | 5 | 774.47 | 586.56 | 6 |
| s_E2 (s) | 8169.94 | 1645.95 | 9 | 5667.59 | 1939.55 | 10 | 173.63 | 158.89 | 7 | 72.52 | 44.64 | 7 | 947.68 | 585.87 | 8 | 491.67 | 332.15 | 11 |
| mx_E2 (s) | 7326.05 | 1644.86 | 9 | 5803.52 | 2158.04 | 8 | 303.86 | 274.17 | 4 | 68.09 | 32.24 | 5 | 1432.46 | 849.95 | 5 | 824.25 | 581.96 | 6 |
| n_sE2 | 1.11 | 0.31 | 9 | 1.30 | 0.30 | 10 | 0.14 | 0.14 | 7 | 0.00 | 0.00 | 7 | 0.25 | 0.16 | 8 | 0.18 | 0.12 | 11 |
| a_sE2 (s) | 8428.09 | 1587.22 | 7 | 4802.68 | 2105.86 | 8 | 1126.18 | NA | 1 | NA | NA | 0 | 3197.12 | 1411.70 | 2 | 2185.45 | 1503.82 | 2 |
| m_sgE2 (s) | 8505.75 | 1547.82 | 7 | 4586.98 | 2146.66 | 8 | 1126.18 | NA | 1 | NA | NA | 0 | 3197.12 | 1411.70 | 2 | 2185.45 | 1503.82 | 2 |
| s_sE2 (s) | 7862.25 | 1719.93 | 9 | 5266.53 | 1956.16 | 10 | 160.88 | 160.88 | 7 | 0.00 | 0.00 | 7 | 799.28 | 587.34 | 8 | 397.35 | 334.91 | 11 |
| t_1Pr (s) | 428.27 | 248.05 | 9 | 380.89 | 112.55 | 10 | 715.24 | 224.31 | 7 | 629.56 | 239.62 | 7 | 817.89 | 307.82 | 8 | 472.68 | 162.69 | 11 |
| d_1Pr (s) | 4764.64 | 2641.67 | 9 | 8871.71 | 2371.04 | 10 | 4892.36 | 1885.44 | 7 | 5275.14 | 3048.54 | 7 | 10525.01 | 3855.33 | 8 | 6898.93 | 2435.22 | 11 |
| t_1E (s) | 3710.37 | 1209.11 | 9 | 5304.17 | 1684.38 | 9 | 5041.51 | 1615.19 | 6 | 8172.30 | 1818.52 | 7 | 2511.57 | 1151.88 | 5 | 10998.41 | 1736.04 | 7 |
| t_C_1E_1Pr (s) | 1340.00 | 515.08 | 9 | 860.90 | 176.13 | 9 | 748.03 | 342.32 | 6 | 2128.63 | 850.38 | 7 | 739.86 | 124.46 | 5 | 860.69 | 194.96 | 7 |
| at_C_1E_Pr (s) | 1302.81 | 332.74 | 8 | 972.67 | 217.13 | 6 | 726.18 | 193.50 | 6 | 2340.73 | 974.89 | 6 | 767.00 | 149.38 | 4 | 1238.33 | 191.50 | 4 |
| mn_C_1E_Pr (s) | 694.00 | 142.28 | 8 | 873.97 | 229.66 | 6 | 552.05 | 156.93 | 6 | 2340.73 | 974.89 | 6 | 697.83 | 184.93 | 4 | 1205.75 | 205.01 | 4 |
| n_bPr_1E | 0.78 | 0.36 | 9 | 0.78 | 0.52 | 9 | 0.83 | 0.48 | 6 | 1.14 | 0.40 | 7 | 0.40 | 0.24 | 5 | 1.43 | 0.65 | 7 |
| n_Pr_1E | 1.33 | 0.47 | 9 | 1.78 | 0.95 | 9 | 1.83 | 0.95 | 6 | 2.43 | 1.00 | 7 | 0.80 | 0.37 | 5 | 4.43 | 1.56 | 7 |
| t_1E12 (s) | 4977.58 | 1517.61 | 9 | 4255.91 | 1352.84 | 8 | 8677.22 | 3252.76 | 4 | 14226.89 | 2564.22 | 7 | 4431.24 | 2127.67 | 5 | 13205.46 | 1922.71 | 6 |
| t_1E2 (s) | 5199.95 | 1481.23 | 9 | 4378.08 | 1338.13 | 8 | 8740.19 | 3253.38 | 4 | 11409.91 | 2578.30 | 5 | 4930.18 | 2000.96 | 5 | 13346.74 | 1873.71 | 6 |
| t_1sE2 (s) | 4676.62 | 1841.94 | 7 | 7409.88 | 2324.45 | 8 | 16118.09 | 5297.77 | 3 | 21455.21 | NA | 1 | 7789.44 | 5631.61 | 3 | 10621.28 | 4792.90 | 2 |
| n_Pr_1E2 | 1.56 | 0.44 | 9 | 1.63 | 1.07 | 8 | 1.25 | 0.25 | 4 | 2.80 | 1.07 | 5 | 0.80 | 0.37 | 5 | 4.67 | 0.71 | 6 |
| n_Pr_1sE2 | 1.29 | 0.52 | 7 | 2.50 | 1.60 | 8 | 2.00 | NA | 1 | NA | NA | 0 | 0.50 | 0.50 | 2 | 5.00 | 3.00 | 2 |
| nPr_1sE2 | 2.00 | 0.79 | 7 | 3.00 | 1.75 | 8 | 2.00 | NA | 1 | NA | NA | 0 | 4.50 | 4.50 | 2 | 0.50 | 0.50 | 2 |
| n_E2_1sE2 | 0.29 | 0.18 | 7 | 0.75 | 0.37 | 8 | 0.00 | NA | 1 | NA | NA | 0 | 1.00 | 1.00 | 2 | 0.50 | 0.50 | 2 |
| t_1E1_1E2 (s) | 1489.58 | 1080.96 | 9 | 950.85 | 674.53 | 9 | 7606.16 | 2957.58 | 6 | 6082.82 | 3090.35 | 7 | 2418.61 | 1854.75 | 5 | 7450.66 | 3007.05 | 9 |
| t_1E1_1sE2 (s) | 8390.86 | 2831.07 | 9 | 8949.96 | 2564.02 | 9 | 18439.45 | 2590.17 | 6 | 20970.44 | 239.62 | 7 | 13235.97 | 4543.07 | 5 | 18745.42 | 1738.83 | 9 |
| s_E1_1sE2 (s) | 529.30 | 165.08 | 7 | 352.46 | 94.06 | 8 | 101.77 | NA | 1 | NA | NA | 0 | 159.91 | 55.06 | 2 | 442.59 | 54.13 | 2 |

| | | | | | | | | | | | | | | | | | | |
|---|---|---|---|---|---|---|---|---|---|---|---|---|---|---|---|---|---|---|
| s_E2_1sE2 (s) | 94.84 | 74.71 | 7 | 157.06 | 97.57 | 8 | 0.00 | NA | 1 | NA | NA | 0 | 185.67 | 185.67 | 2 | 68.42 | 68.42 | 2 |
| a_E2_1sE2 (s) | 331.94 | 196.27 | 2 | 209.42 | 92.25 | 3 | NA | NA | 0 | NA | NA | 0 | 185.67 | NA | 1 | 136.83 | NA | 1 |
| rel_E2_C | 2.29 | 0.74 | 9 | 2.49 | 1.96 | 9 | 0.02 | 0.02 | 6 | 0.02 | 0.01 | 7 | 0.16 | 0.07 | 5 | 0.16 | 0.13 | 7 |
| rel_E1_allE | 0.13 | 0.05 | 9 | 0.11 | 0.03 | 9 | 0.43 | 0.18 | 6 | 0.45 | 0.13 | 7 | 0.41 | 0.14 | 5 | 0.37 | 0.11 | 7 |
| n_frE1_n_E12 | 1.06 | 0.06 | 9 | 0.92 | 0.12 | 9 | 0.83 | 0.31 | 6 | 0.71 | 0.18 | 7 | 1.20 | 0.20 | 5 | 0.86 | 0.14 | 7 |
| rel_E2_1E2 | 0.49 | 0.09 | 9 | 0.39 | 0.11 | 8 | 0.02 | 0.02 | 4 | 0.01 | 0.00 | 5 | 0.08 | 0.04 | 5 | 0.08 | 0.03 | 6 |
| n_pd | 47.56 | 12.28 | 9 | 64.00 | 12.84 | 10 | 77.29 | 17.44 | 7 | 66.14 | 16.11 | 7 | 39.50 | 11.98 | 8 | 57.91 | 10.47 | 11 |
| a_pd | 5.10 | 0.36 | 9 | 4.88 | 0.40 | 9 | 4.70 | 0.21 | 7 | 5.01 | 0.36 | 7 | 4.61 | 0.24 | 8 | 4.79 | 0.28 | 11 |
| m_pd (s) | 4.87 | 0.37 | 9 | 4.79 | 0.25 | 9 | 4.65 | 0.26 | 7 | 4.92 | 0.32 | 7 | 4.66 | 0.23 | 8 | 4.60 | 0.23 | 11 |
| s_pd (s) | 240.92 | 60.38 | 9 | 312.95 | 62.18 | 10 | 369.62 | 88.34 | 7 | 316.01 | 70.40 | 7 | 173.89 | 47.56 | 8 | 271.88 | 48.55 | 11 |
| a_pd_II_1 (s) | 1.94 | 0.26 | 5 | 1.80 | 0.15 | 4 | 2.30 | 0.75 | 4 | 1.72 | 0.06 | 4 | 1.91 | NA | 1 | 2.04 | 0.27 | 4 |
| m_pd_II_1 (s) | 1.91 | 0.27 | 5 | 1.81 | 0.15 | 4 | 2.28 | 0.76 | 4 | 1.73 | 0.05 | 4 | 1.90 | NA | 1 | 2.05 | 0.27 | 4 |
| s_pd_II_1 (s) | 4.00 | 1.82 | 9 | 5.78 | 3.58 | 10 | 10.51 | 4.58 | 7 | 3.32 | 2.23 | 7 | 2.15 | 2.15 | 8 | 6.31 | 3.84 | 11 |
| a_pd_II_2 (s) | 0.86 | 0.05 | 5 | 0.78 | 0.05 | 4 | 1.42 | 0.44 | 4 | 0.98 | 0.07 | 4 | 0.93 | NA | 1 | 0.93 | 0.05 | 3 |
| m_pd_II_2 (s) | 0.88 | 0.07 | 5 | 0.79 | 0.06 | 4 | 1.41 | 0.44 | 4 | 0.97 | 0.07 | 4 | 0.88 | NA | 1 | 0.91 | 0.09 | 3 |
| s_pd_II_2 (s) | 2.03 | 0.96 | 9 | 2.58 | 1.66 | 10 | 6.71 | 2.98 | 7 | 2.16 | 1.41 | 7 | 1.05 | 1.05 | 8 | 3.05 | 1.91 | 11 |
| a_pd_II_3 (s) | 2.69 | 0.35 | 5 | 2.66 | 0.28 | 4 | 3.22 | 0.71 | 4 | 2.62 | 0.42 | 4 | 2.33 | NA | 1 | 2.73 | 0.32 | 4 |
| m_pd_II_3 (s) | 2.62 | 0.40 | 5 | 2.53 | 0.29 | 4 | 3.21 | 0.73 | 4 | 2.60 | 0.43 | 4 | 2.13 | NA | 1 | 2.75 | 0.30 | 4 |
| s_pd_II_3 (s) | 6.15 | 3.05 | 9 | 8.39 | 5.25 | 10 | 16.64 | 7.53 | 7 | 5.05 | 3.05 | 7 | 2.33 | 2.33 | 8 | 10.14 | 6.96 | 11 |
| t_1pd (s) | 453.12 | 172.94 | 9 | 393.09 | 225.29 | 9 | 288.73 | 115.97 | 7 | 582.69 | 435.60 | 7 | 543.51 | 259.67 | 8 | 334.56 | 111.91 | 11 |
| t_1pd_1pr (s) | 155.93 | 39.44 | 9 | 157.23 | 50.72 | 9 | 149.63 | 66.28 | 7 | 124.38 | 50.27 | 7 | 260.33 | 128.24 | 8 | 297.17 | 112.47 | 11 |
| at_1pd_Pr (s) | 232.79 | 104.55 | 9 | 120.81 | 17.87 | 9 | 151.12 | 36.32 | 7 | 156.72 | 43.34 | 7 | 259.87 | 128.42 | 8 | 155.19 | 20.60 | 11 |
| mt_1pd_Pr (s) | 128.88 | 38.29 | 9 | 108.11 | 18.86 | 9 | 69.13 | 10.75 | 7 | 140.71 | 47.39 | 7 | 255.48 | 129.16 | 8 | 126.24 | 18.98 | 11 |
| mnt_1pd_1Pr (s) | 63.56 | 32.06 | 9 | 53.37 | 19.47 | 9 | 29.40 | 7.87 | 7 | 112.89 | 52.52 | 7 | 242.76 | 131.97 | 8 | 73.22 | 22.02 | 11 |
| n_pd_minC_pd | 0.50 | 0.08 | 9 | 0.47 | 0.08 | 10 | 0.77 | 0.14 | 7 | 0.68 | 0.12 | 7 | 0.70 | 0.18 | 8 | 0.52 | 0.08 | 11 |
| n_pd_minC | 0.47 | 0.08 | 9 | 0.44 | 0.07 | 10 | 0.70 | 0.13 | 7 | 0.64 | 0.11 | 7 | 0.65 | 0.16 | 8 | 0.49 | 0.07 | 11 |
| n_pd_1Pr | 15.89 | 4.90 | 9 | 23.60 | 5.89 | 10 | 27.14 | 8.18 | 7 | 21.14 | 4.61 | 7 | 23.38 | 5.08 | 8 | 16.00 | 5.10 | 11 |
| rel_Prob_pd | 1.00 | 0.00 | 9 | 0.90 | 0.10 | 10 | 1.00 | 0.00 | 7 | 1.00 | 0.00 | 7 | 1.00 | 0.00 | 8 | 1.00 | 0.00 | 11 |
| n_Pr_1pd | 0.56 | 0.18 | 9 | 0.60 | 0.16 | 10 | 0.71 | 0.18 | 7 | 0.71 | 0.18 | 7 | 0.25 | 0.16 | 8 | 0.64 | 0.15 | 11 |
| d_1pd (s) | 5.53 | 1.02 | 9 | 3.32 | 0.64 | 10 | 5.11 | 1.12 | 7 | 4.80 | 0.82 | 7 | 3.64 | 0.67 | 8 | 6.44 | 1.07 | 11 |
| d_2pd (s) | 6.27 | 0.66 | 9 | 3.93 | 0.82 | 10 | 5.48 | 0.94 | 7 | 4.04 | 0.67 | 7 | 4.41 | 0.62 | 8 | 3.95 | 0.67 | 11 |
| d_pd5 (s) | 5.25 | 0.44 | 9 | 3.97 | 0.65 | 9 | 5.34 | 0.59 | 7 | 4.67 | 0.61 | 7 | 4.01 | 0.46 | 8 | 4.63 | 0.60 | 11 |
| s_pdII_3_5pd (s) | 0.37 | 0.37 | 9 | 0.58 | 0.58 | 10 | 3.95 | 2.23 | 7 | 0.45 | 0.45 | 7 | 0.00 | 0.00 | 8 | 0.00 | 0.00 | 11 |

Standard EPG phase codes: np = non-probing, Pr = tissue probe by an aphid stylet, bPr = brief probe (< 3 mins), C = pathway phase (stylet interaction with upper epidermal/mesophyll tissue), pd = potential drop (intracellular puncture by aphid stylet), F = derailed stylet mechanics, G = xylem ingestion, E1 = salivation into the phloem, E1e = salivation into non-phloem tissue, E2 = phloem ingestion, sE2 = sustained phloem ingestion (>10 minutes). Specific parameter codes used by Schliephake et al., 2013: n = number of observed events, s = total, m = median, a = average, d = duration, mx = maximum, mn = minimum, sg = the longest single event, fr = fraction, t = time passed before a parameter was observed in the recording, at = average time, mt = median time, mnt = minimum time, min = per minute, rel_E2_C = ingestion:pathway ratio, rel_E1_allE = E1 index, rel_E2_1E2 = E2 index, n_frE1_n_E12 = phloem phase fractioning. Description of the NA values: in the mean column NA indicates that this parameter was not observed in any replicate for the respective treatment, in the SE column NA indicates that the standard error could not be calculated (i.e. no observations or parameter was only observed in one replicate).

**Table S6:** Statistical results of plant physiological responses. Physiological parameters were analysed in response to the imposed water treatment, plant type, aphid presence, and all interactions. Bold text indicates significant p-values.

| Explanatory Variable | Plant Physiological Parameter | | | | | |
|---|---|---|---|---|---|---|
| | **Development Time** | **Mean Stomatal Conductance** | **Shoot Dry Mass** | **Root Dry Mass** | **Root Length** | **Root:Shoot** |
| Water Treatment | $X^2_2$ = 167.56; **p = <0.001** | $X^2_2$ = 85.39; p = <0.001 | $F_{2, 135}$ = 190.80; **p = <0.001** | $F_{2, 143}$ = 39.26; **p = <0.001** | $F_{2, 141}$ = 45.71; **p = <0.001** | $F_{2, 140}$ = 49.89; **p = <0.001** |
| Plant Type | $X^2_1$ = 1.49; p = 0.222 | $X^2_1$ = 31.67; **p = <0.001** | $F_{1, 135}$ = 13.35; **p = <0.001** | $F_{1,142}$ = 1.45; p = 0.230 | $F_{1, 141}$ = 8.72; **p = 0.003** | $F_{1, 139}$ = 5.43; **p = 0.021** |
| Aphid Presence | N/A | N/A | $F_{1, 135}$ = 1.05; p = 0.307 | $F_{1, 140}$ = 1.01; p = 0.314 | $F_{1, 140}$ = 2.26; p = 0.134 | $F_{1, 138}$ = 0.58; p = 0.447 |
| Water Treatment x Plant Type | $X^2_2$ =0.71; p = 0.698 | $X^2_2$ =1.00; p = 0.606 | $F_{2, 135}$ = 2.62; p = 0.075 | $F_{2, 137}$ = 1.05; p = 0.351 | $F_{2, 137}$ = 0.71; p = 0.489 | $F_{2, 143}$ = 0.24; p = 0.782 |
| Water Treatment x Aphid Presence | N/A | N/A | $F_{2, 135}$ = 0.13; p = 0.871 | $F_{2, 135}$ = 0.08; p = 0.922 | $F_{2, 135}$ = 0.16; p = 0.848 | $F_{1, 135}$ = 0.29; p = 0.777 |
| Plant Type x Aphid Presence | N/A | N/A | $F_{1, 135}$ = 3.56; p = 0.061 | $F_{1, 137}$ = 0.12; p = 0.719 | $F_{1, 139}$ = 2.93; p = 0.088 | $F_{1, 137}$ = 0.68; p = 0.409 |
| Water Treatment x Plant Type x Aphid Presence | N/A | N/A | $F_{2, 135}$ = 0.22; p = 0.802 | $F_{2, 132}$ = 0.26; p = 0.771 | $F_{2, 133}$ = 0.76; p = 0.468 | $F_{2, 141}$ = 0.61; p = 0.433 |

N/A: These parameters were measured before aphid infestation so no statistical results are available for aphid presence.

**Table S7:** Statistical results for the measured aphid fitness and EPG parameters. Parameters were analysed in response to the imposed water treatment, plant type, and the interaction. Bold text indicates significant p-values.

| **Explanatory Variable** | **Aphid Fitness Parameter** | | **EPG parameter** |
|---|---|---|---|
| | **Nymph Dry Mass** | **Cumulative Aphid Fecundity** | **Multivariate analysis of all parameters** |
| Water Treatment | $F_{2,64}$ = 15.66; **p = <0.001** | $F_{2,61}$ = 18.83; **p = <0.001** | $F_{1,46}$ = 6.42; **p = 0.001** |
| Plant Type | $F_{1,63}$ = 0.09; p = 0.757 | $F_{1,61}$ = 0.61; p = 0.435 | $F_{2,46}$ = 1.24; p = 0.302 |
| Water Treatment x Plant Type | $F_{2,61}$ = 2.29; p = 0.109 | $F_{2,61}$ = 3.12; p = 0.050 | $F_{2,46}$ = 1.17; p = 0.323 |
